# Immunosuppression with Cyclosporine versus Tacrolimus shows distinctive nephrotoxicity profiles within renal compartments

**DOI:** 10.1101/2023.04.05.535688

**Authors:** Hasan Demirci, Suncica Popovic, Carsten Dittmayer, Duygu Elif Yilmaz, Ismail Amr El-Shimy, Michael Mülleder, Christian Hinze, Pontus B. Persson, Kerim Mutig, Sebastian Bachmann

## Abstract

Calcineurin inhibitors (CNI) are the backbone for immunosuppression after solid organ transplantation. Although successful in preventing kidney transplant rejection, their nephrotoxic side effects notoriously contribute to allograft injury despite attempts to optimize their application, often with additional medications. Complex renal parenchymal damage occurs for cyclosporine A (CsA) as well as for the currently favoured tacrolimus (Tac). To test for distinct CsA and Tac damaging patterns, we combined multiomics analysis with histopathology from rat kidneys exposed to continuous CNI delivery. Damage forms varied strikingly. Both drugs caused significant albeit differential damage in vasculature and nephron. The glomerular filtration barrier was more affected by Tac than by CsA, showing prominent deteriorations in pore endothelium and podocytes along with impaired VEGF/VEGFR2 signaling and podocyte-specific gene expression. By contrast, proximal tubule epithelia were more severely affected by CsA than by Tac, revealing lysosomal dysfunction and enhanced apoptosis along with impaired proteostasis and oxidative stress. We conclude that pathogenetic alterations in renal microenvironments are specific for either treatment. Should this translate to the clinical setting, CNI choice should reflect individual risk factors for renal vasculature and tubular epithelia. As a step in this direction, we share products identified from multiomics for differential pathognomonic biomarkers.

**Translational Statement:** Calcineurin inhibitors (CNI) are first-choice immunosuppressive agents. Their nephrotoxic side effects may often limit their use. Tacrolimus is currently preferred to cyclosporine although its superiority remains unclear. Within the nephron, damage to the filtration barrier is greater for tacrolimus, whereas cyclosporine side effects locate more to the proximal tubular epithelium when compared in our rodent model. We identify the distinctive location and nature of damage by both drugs and unravel involved mechanisms. By detecting differential protein signatures we make available pathognomonic biomarkers for renal allograft health under CNI treatment.

## Introduction

Calcineurin inhibitors (CNI) such as cyclosporine A (CsA) and tacrolimus (Tac, FK506) have become the basis for immunosuppressive treatment in organ transplant recipients and patients with autoimmune diseases over the past four decades. Today, over 90% of kidney transplant recipients in Western countries are maintained on CNI-containing immunosuppressive regimens. Apart from a satisfactory outcome at short term, chronic use of CNI causes nephrotoxicity (NTX) in a significant proportion of renal and nonrenal transplant recipients, affecting renal allograft function^1–3^. CNI elimination or substituting with other immunosuppressant regimens are explored, but CNI alternatives are widely considered inferior^1, 4, 5^. CNI-NTX with decreasing allograft function may root from chronic hypoperfusion related to hyaline arteriolopathy with glomerular scarring, direct tubular toxicity causing vacuolation, atrophy, and striped fibrosis. Their interplay leads to interstitial fibrosis/tubular atrophy (IF/TA). Immunological damage and comorbidities unrelated to CNI add to potential CNI injury. Thus, whether certain pathological landmarks occur specifically for CsA or Tac remains to be determined^2, 6–8^. Should the two CNI substance groups pose a threat to unlike kidney structures, CNI choice should reflect the patient’s history and comorbidities. Calcineurin is a calcium/calmodulin-dependent serine-threonine phosphatase and notable for its key role in T cell function. Upon activation, the catalytic calcineurin subunit (CnA) associates with the regulatory subunit (CnB) to dephosphorylate and thereby activate transcription factors of the NFAT (nuclear factor of activated T-cells) family. NFAT induce expression of key genes for T-lymphocyte differentiation^9, 10^. Calcineurin and NFAT isoforms are, however, not T-cell specific and may encompass interactions with other substrates targeting proteins and regulators of calcineurin activity beyond immunology. CNI-NTX can involve vasoconstriction, hypertension, tissue hypoxia, oxidative stress, anemia, hyperkalemia, metabolic acidosis, dysregulation of major endo- and paracrine systems, and metabolic stress^11–15^. Although widely assumed that CsA and Tac produce identical lesions^7^, this view is not unopposed. CsA may cause greater nephrotoxicity than Tac^3, 16, 17^. Both CNI inhibit calcineurin activity by forming complexes with distinct members of the immunophilin family, i.e. cyclophilins for CsA and FKBP12 for Tac. These distinct actions may explain the specific toxicity profiles of various CNIs. Distinctively, CsA affects proteostasis, since the chaperone activity of cyclophilins interacts with protein maturation^16^. Studies comparing CsA vs. Tac in cell culture consequently show stronger association of CsA with ER stress and the unfolded protein response (UPR), leading to accumulation of misfolded protein and apoptotic cell death, whereas Tac did not^17–20^. CsA-NTX may further involve altered JAK/STAT signaling and salt-sensitive mitochondrial dysfunction^21^. In search of potential nephroprotection from CsA, transcription factor EB (TFEB)-mediated autophagy flux and lysosomal dysfunction is discussed, owing to CsA-induced mTOR overactivity ^22^. The current preference to provide Tac rather than CsA^3^ (except for new onset of diabetes cases^1^ and specific combination therapies) requires substantiation. By understanding cellular mechanisms of NTX for both substance families, better-tailored treatment protocols may arise to reduce CNI adverse effects in the individual patient. We test whether CsA and Tac exposure, before reaching irreversible damage, affect renal compartments differently, based on their different pathogenic pathways. Our results identify distinctive mechanisms and related biomarkers in Tac vs. CsA groups serving to link individual pathways to nephroprotective modifications in patient immunosuppressive protocols.

## Short Methods

See online supplementary methods for further detail.

### Animals

As males are more prone to kidney injury, adult male Wistar rats received CsA (Sandimmun, Novartis; target plasma trough level 3 µg/ml), Tac (Selleckchem; target plasma trough level 3.5 ng/ml) or respective vehicle via subcutaneously implanted osmotic minipumps (Alzet, 2ML4) for 28 days. Rats were placed in metabolic cages to collect urine. Blood samples and kidneys were taken for biochemical evaluation. Animals were perfusion-fixed via the abdominal aorta and kidneys prepared as described^23^.

### Microscopy

For histology and immunohistochemistry (IHC), 4 μm-thick sections were stained with PAS, Masson’s Trichrome, or Sirius Red. For IHC, epitope retrieval and staining protocols were used as described^14^. For electron microscopy, a Gemini 300 field emission scanning electron microscope (FESEM, Zeiss) equipped with a STEM detector was employed for large-scale digitization^23^. High-resolution TIF files were inspected with QuPath and transferred to a repository (www.nanotomy.org) for open access pan-and-zoom analysis.

### Morphometry

Tubulo-interstitial fibrosis, arteriolar dimensions, endothelial and epithelial signals were evaluated by histology and IHC from multiple samples and optical fields using ImageJ. Glomerular endothelial pore density and fenestration density of peritubular capillaries were quantified in TEM images. Epithelial lysosomes were counted in Richardson’s stained semithin plastic sections. For filtration slit density (FSD) measurements, podocyte exact morphology measurement procedure (PEMP; NIPOKA) was applied using 3D-structured illumination microscopy (3D-SIM)^24, 25^.

### Multiomics analysis

RNA-seq of kidney RNA samples (n=4 to 6 per group) was performed (Novogene). Global proteomics and phosphoproteomics from kidney protein lysates were done by the Charité and Max Delbrück Center core facilities; data were deposited in the NCBI’s Gene Expression Omnibus repository (GSE225215), ProteomeXchange Consortium (PXD038841, PXD038546).

## Results

### Physiologic Parameters

CNI delivered for 4 weeks by osmotic minipumps resulted in plasma trough levels that correspond to established data (3.04 µg/ml in the CsA and 3.42 ng/ml in the Tac groups)^26, 27^. CNI-induced kidney injury was assessed by reduced creatinine clearance, elevated blood urea nitrogen (BUN) and cystatin C levels, and reduced fractional Na^+^ excretion (Table 1, Fig. 1).

**Table 1.** Physiologic parameters, numerical values. Urine and blood parameters were obtained from rats after 4 weeks of treatment with vehicle (Veh), cyclosporine A (CsA), or tacrolimus (Tac).

| Parameter | Veh (n=5) | CsA (n=5) | P value | Veh (n=6) | Tac (n=6) | P value |
| --- | --- | --- | --- | --- | --- | --- |
| **Serum** |  |  |  |  |  |  |
| Na<sup>+</sup> (mmol/l) | 139 ± 5 | 142 ± 1 | 0.31 | 144 ± 2 | 143 ± 1 | 0.282 |
| Cl<sup>-</sup> (mmol/l) | 97 ± 3 | 92 ± 4 | 0.039* | 103 ± 1 | 101 ± 1 | 0.002** |
| K<sup>+</sup> (mmol/l) | 4.25 ± 0.77 | 4.50 ± 0.45 | 0.547 | 4.95 ± 0.80 | 4.70 ± 0.53 | 0.546 |
| Ca<sup>+2</sup> (mg/dl) | 10.4 ± 0.8 | 9.7 ± 0.9 | 0.238 | 9.7 ± 0.4 | 10.1 ± 0.3 | 0.095 |
| Phosphate (mg/dl) | 8.09 ± 1.04 | 8.09 ± 0.49 | 0.994 | 9.06 ± 0.68 | 8.41 ± 0.54 | 0.094 |
| Uric acid (mg/dl) | 0.86 ± 0.85 | 0.92 ± 0.39 | 0.887 | 1.48 ± 0.97 | 1.13 ± 0.33 | 0.447 |
| Lactate (mg/dl) | 35.9 ± 18.3 | 37.5 ± 12.9 | 0.874 | 34.2 ± 6.9 | 32.8 ± 4.8 | 0.697 |
| Glucose (mg/dl) | 147 ± 14 | 179 ± 3 | 0.034* | 151 ± 13 | 172 ± 10 | 0.019* |
| Urea (mg/dl) | 34.2 ± 2.8 | 40.0 ± 4.8 | 0.06 | 29.7 ± 3.5 | 8.7 ± 5.8 | 0.011* |
| Creatinine (mg/dl) | 0.31 ± 0.02 | 0.33 ± 0.02 | 0.216 | 0.29 ± 0.01 | 0.32 ± 0.01 | 0.001*** |
| Cystatin C (mg/l) | 0.40 ± 0.07 | 0.53 ± 0.04 | 0.01** | 0.38 ± 0.06 | 0.49 ± 0.07 | 0.02* |
| Trough level (ng/ml) | 0 | 3040 ± 126 | - | 0 | 3.42 ± 0.80 | - |
| Parameter | Veh (n=5) | CsA (n=5) | P value | Veh (n=6) | Tac (n=6) | P value |
| **Urine** |  |  |  |  |  |  |
| Na<sup>+</sup> (mmol/kg) | 4.53 ± 0.72 | 2.29 ± 0.88 | 0.001*** | 6.11 ± 0.94 | 3.54 ± 1.08 | 0.001*** |
| Cl<sup>-</sup> (mmol/kg) | 7.34 ± 0.81 | 6.86 ± 1.15 | 0.44 | 9.93 ± 1.03 | 12.14 ± 1.38 | 0.01** |
| K<sup>+</sup> (mmol/kg) | 10.19 ± 0.32 | 9.43 ± 0.91 | 0.1 | 12.04 ± 0.91 | 15.10 ± 1.34 | 0.001*** |
| Phosphate (mg/kg) | 8.8 ± 8.4 | 43.3 ± 8.9 | 0.001*** | 10.1 ± 4.7 | 60.0 ± 12.2 | 0.0001**** |
| Glucose (mg/kg) | 11 ± 10 | 562 ± 588 | 0.07 | 20 ± 9 | 1204 ± 1444 | 0.1 |
| Urea (mg/kg) | 1470 ± 93 | 1380 ± 209 | 0.39 | 1810 ± 221 | 2310 ± 255 | 0.005** |
| Creatinine (mg/kg) | 24.1 ± 3.0 | 20.1 ± 1.9 | 0.04* | 30.1 ± 1.3 | 30.5 ± 1.7 | 0.67 |
| Albumin (g/kg) | 0.06 ± 0.03 | 0.14 ± 0.06 | 0.02* | 0.22 ± 0.03 | 0.19 ± 0.11 | 0.59 |
| Urine flow (ml/24h) | 23.0 ± 6.0 | 14.3 ± 3.4 | 0.03* | 16.0 ± 3.0 | 21.3 ± 5.5 | 0.07 |
| Osmolality (mOsm/kg) | 1270 ± 364 | 1660 ± 268 | 0.08 | 1630 ± 209 | 1490 ± 214 | 0.3 |

**Table 1. Physiologic parameters, numerical values.**
| Parameter | Veh (n=5) | CsA (n=5) | *P* value | Veh (n=6) | Tac (n=6) | *P* value |
| --- | --- | --- | --- | --- | --- | --- |
| **Serum** |  |  |  |  |  |  |
| Na<sup>+</sup> [mmol/l] | 139 ± 5 | 142 ± 1 | 0.31 | 144 ± 2 | 143 ± 1 | 0.282 |
| Cl<sup>-</sup> [mmol/l] | 97 ± 3 | 92 ± 4 | 0.039* | 103 ± 1 | 101 ± 1 | 0.002** |
| K<sup>+</sup> [mmol/l] | 4.25 ± 0.77 | 4.50 ± 0.45 | 0.547 | 4.95 ± 0.80 | 4.70 ± 0.53 | 0.546 |
| Ca<sup>+2</sup> [mg/dl] | 10.4 ± 0.8 | 9.7 ± 0.9 | 0.238 | 9.7 ± 0.4 | 10.1 ± 0.3 | 0.095 |
| Phosphate [mg/dl] | 8.09 ± 1.04 | 8.09 ± 0.49 | 0.994 | 9.06 ± 0.68 | 8.41 ± 0.54 | 0.094 |
| Uric acid [mg/dl] | 0.86 ± 0.85 | 0.92 ± 0.39 | 0.887 | 1.48 ± 0.97 | 1.13 ± 0.33 | 0.447 |
| Lactate [mg/dl] | 35.9 ± 18.3 | 37.5 ± 12.9 | 0.874 | 34.2 ± 6.9 | 32.8 ± 4.8 | 0.697 |
| Glucose [mg/dl] | 147 ± 14 | 179 ± 3 | 0.034* | 151 ± 13 | 172 ± 10 | 0.019* |
| Urea [mg/dl] | 34.2 ± 2.8 | 40.0 ± 4.8 | 0.06 | 29.7 ± 3.5 | 8.7 ± 5.8 | 0.011* |
| Creatinine [mg/dl] | 0.31 ± 0.02 | 0.33 ± 0.02 | 0.216 | 0.29 ± 0.01 | 0.32 ± 0.01 | 0.001*** |
| Cystatin C [mg/l] | 0.40 ± 0.07 | 0.53 ± 0.04 | 0.01** | 0.38 ± 0.06 | 0.49 ± 0.07 | 0.02* |
| Trough level [ng/ml] | 0 | 3040 ± 126 | - | 0 | 3.42 ± 0.80 | - |
| **Urine** |  |  |  |  |  |  |
| Na<sup>+</sup> [mmol/kg] | 4.53 ± 0.72 | 2.29 ± 0.88 | 0.001*** | 6.11 ± 0.94 | 3.54 ± 1.08 | 0.001*** |
| Cl<sup>-</sup> [mmol/kg] | 7.34 ± 0.81 | 6.86 ± 1.15 | 0.44 | 9.93 ± 1.03 | 12.14 ± 1.38 | 0.01** |
| K<sup>+</sup> [mmol/kg] | 10.19 ± 0.32 | 9.43 ± 0.91 | 0.1 | 12.04 ± 0.91 | 15.10 ± 1.34 | 0.001*** |
| Phosphate [mg/kg] | 8.8 ± 8.4 | 43.3 ± 8.9 | 0.001*** | 10.1 ± 4.7 | 60.0 ± 12.2 | 0.0001**** |
| Glucose [mg/kg] | 11 ± 10 | 562 ± 588 | 0.07 | 20 ± 9 | 1204 ± 1444 | 0.1 |
| Urea [mg/kg] | 1470 ± 93 | 1380 ± 209 | 0.39 | 1810 ± 221 | 2310 ± 255 | 0.005** |
| Creatinine [mg/kg] | 24.1 ± 3.0 | 20.1 ± 1.9 | 0.04* | 30.1 ± 1.3 | 30.5 ± 1.7 | 0.67 |
| Albumin [g/kg] | 0.06 ± 0.03 | 0.14 ± 0.06 | 0.02* | 0.22 ± 0.03 | 0.19 ± 0.11 | 0.59 |
| Urine flow [ml/24h] | 23.0 ± 6.0 | 14.3 ± 3.4 | 0.03* | 16.0 ± 3.0 | 21.3 ± 5.5 | 0.07 |
| Osmolality [mOsm/kg] | 1270 ± 364 | 1660 ± 268 | 0.08 | 1630 ± 209 | 1490 ± 214 | 0.3 |

Values, normalized for individual body weights, are means ± SD; \**P*<0.05, \*\**P*<0.01,

**Figure 1.**
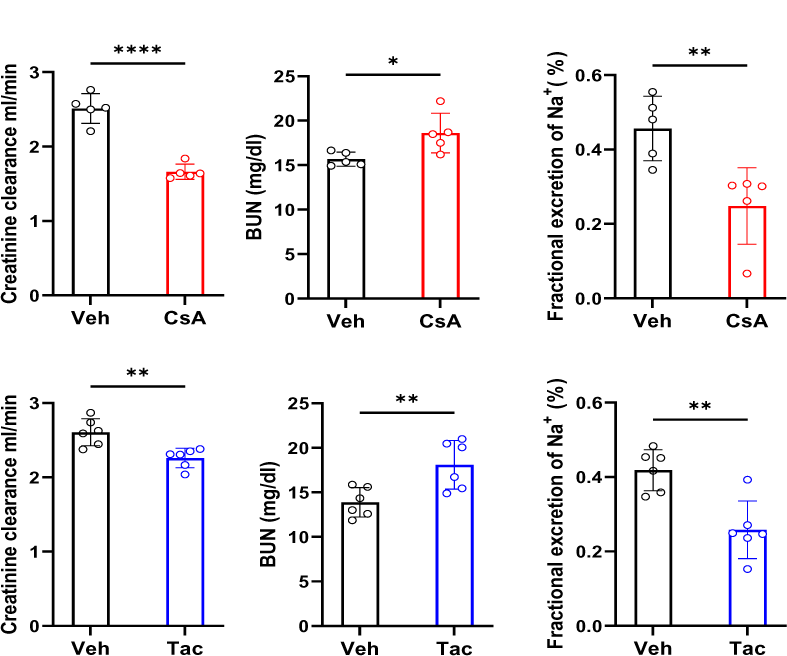
Physiologic parameters, graphs. Creatinine clearance, blood urea nitrogen (BUN), fractional sodium excretion, and urine sodium concentration were obtained from rats after 4 weeks of vehicle (Veh)-, cyclosporine A (CsA)- or tacrolimus (Tac)-treatment. Values, normalized for individual body weights, are means from n= 5 to 6 rats ± SD; * *P*<0.05, \*\**P*<0.01; \*\*\*\**P*<0.0001. Statistical tests were performed using unpaired Student’s *t* test. Table 1.

### Overview Pathology

Substantial pathological changes took place in CNI-recipient kidneys, which ranged between mild onset stages and nephron destruction, reflecting CNI nephropathy. Mild-to-severe glomerular and tubulo-interstitial degenerative alterations were observed within numerous scattered, circumscribed foci of the cortex, while other areas appeared normal. These foci were frequent in the subcapsular cortex but also extending to deeper parts of cortex and outer medulla, initiating striped fibrosis (PAS staining; Fig. 2a-c). Adjacent glomeruli often displayed a retracted, albeit fully perfused phenotype. Interstitium of the foci was loose and meshwork-like in CsA, but dense and cell-enriched in Tac; fibrosis was identified by Trichrome and Sirius red staining (Fig. 2d-j). α-SMA-immunoreactive signal suggested intensive profibrotic myofibroblast transdifferentiation and pericyte activation with pericapsular signals stronger in CsA than in Tac despite similar mean extension of signal (Suppl. Fig. S1a-d). Focal outer medullary interstitial α-SMA signal was equally enhanced in Tac and CsA (Suppl. Fig. S1e-g). In the inner medulla, changes in α-SMA were minimal.

**Figure 2.**
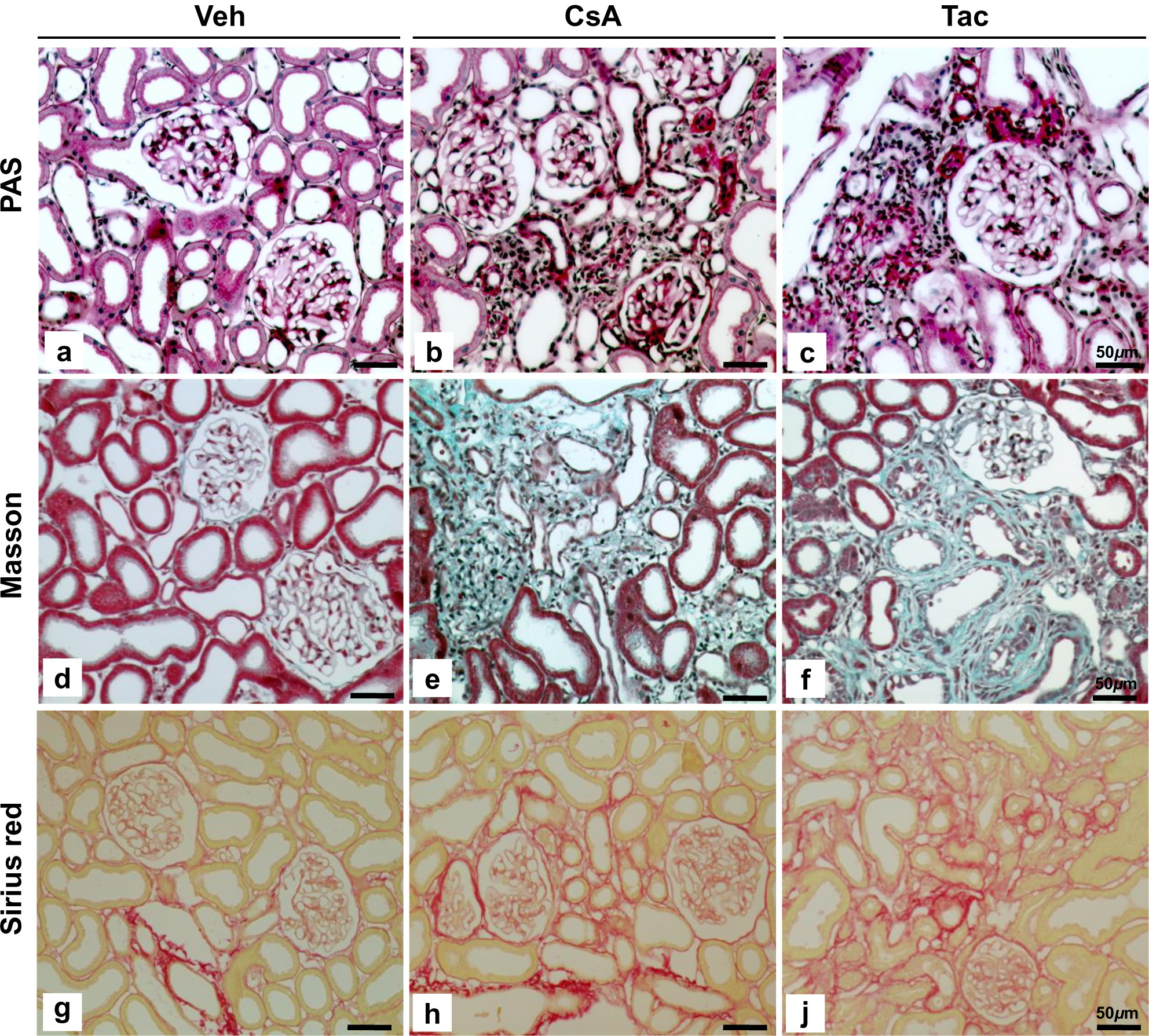
Histological overview. (a-c) Representative overviews of fibrotic foci in cyclosporine A (CsA) and tacrolimus (Tac); periodic acid-Schiff (PAS). Typically, foci consist of 5 to 20 tubular profiles, but also larger areas measuring up to 150.000 µm^2^ were encountered. Note thickening of peritubular basement membrane staining in (b,c). Mean proportional area occupied by foci was 4.0±1.7% for CsA and 4.5±1.3% in Tac (difference not significant; unpaired Student’s *t*-test). (d-f) Green signal characterizes fibrotic foci in CsA and Tac (Masson’s trichrome staining). (g-j) Sirius red staining indicates collagen deposits in fibrotic foci of CsA and Tac. Bars indicate magnification.

Typical cellular infiltrates were scarce, since anti-CD45 immunofluorescence indicated only mild infiltration in fibrotic foci of CsA and Tac samples (Suppl. Fig. S1h-k).

### Renal Vasculature

Interlobular arteries, showed mild increases in wall thickness upon CNI with occasional hyalinic or necrotizing subintimal or media inclusions in CsA as opposed to significant increases in intermyocyte matrix and adventitial collagen as well as myocyte thinning in Tac (Suppl. Fig. S2a-c). Glomerular afferent arterioles showed PAS-positive wall inclusions in CsA and increased wall-to-lumen ratio in Tac. Endothelial CD31 (PECAM-1) expression showed mildly increased signal in CsA and Tac. Ultrastructurally, myocytes were particularly rich in contractile apparatus of single-layered media in CsA, but flattened and layered in Tac (Suppl. Fig. S2d-n). Terminal afferent arterioles regularly displayed hypergranularity of the juxtaglomerular granular cells with both treatments, with renin signal intensities increased 7-fold in CsA and 8.8-fold in Tac compared to controls. Specifically in the Tac group layers of granular cells featured narrowing of the afferent arteriole focally at the glomerular entry (Suppl. Fig. S3a-e). TEM detail of granular cells was inconspicous in CsA but showed glycogen accumulations, necrosis, or hyaline formations in Tac (Suppl. Fig. S3f-h). Interlobular veins and ascending vasa recta viewed by SEM were normal in CsA but strikingly revealed partial or total loss of fenestrae in Tac. Focally adhering CD45-positive leukocytes were seen in both CNI groups, but more frequently in Tac (Suppl. Fig. S4).

### Glomeruli

Glomeruli with mild sclerosis and/or retraction were found both groups in association with fibrotic foci. Capillaries were narrowed, and the mesangium mildly expanded (Fig. 3a). Degenerative changes of the glomerular tuft varied with the degree of retraction and podocyte effacement. By SEM, partial podocyte effacement was commonly found in CsA, whereas in Tac, more extensive or generalized effacement with major podocyte degeneration was detected (Fig. 3b). TEM confirmed higher effacement in Tac than in CsA (Fig. 3c). FSD analysis showed that besides areas of normal configuration of the slit diaphragm sites of significant effacement were detected in both groups. Significant FSD reduction was found in both CNI groups (Fig. 3d). Ultrastructure of the capsule revealed an activated, thickened parietal epithelium with occasional vacuolization, layering of the basement membrane and adjacent hyaline or granular matrix formations. Multiple synechiae with podocytes were commonly observed in CsA; changes in Tac were comparable (Suppl. Fig. S5a-f). CD44 immunostaining specified its activated state sharp increases in parietal epithelial signal in CsA and Tac; pericapsular interstitium was mildly (CsA) or substantially (Tac) infiltrated by CD45-positive leukocytes (Suppl. Fig. S5g-h). Endothelial fenestration of the glomerular capillaries showed conspicous reductions in pore density by SEM which were markedly stronger in Tac than in CsA (Fig. 4a-d). These changes were confirmed by TEM, revealing also substantial condensation of endothelial cells chiefly in Tac (Fig. 4e-k). Areas of reduced pore formation were commonly facing those of podocyte effacement. Reductions in pore density were numerically confirmed (Fig. 4l). Changes of the tuft structures were further paralleled by diminished Wilms tumor 1 protein (WT1) immunoreactivity (Fig. 5a-c). Cell death-associated DNA fragmentation of the tuft by TUNEL immunofluorescence was highly increased in the Tac, but less so in the CsA group; contrastingly, parietal epithelial cells showed a stronger signal in CsA than in Tac (Fig. 5d-f). These observations were confirmed by numerical quantification (Fig. 5g-j).

**Figure 3.**
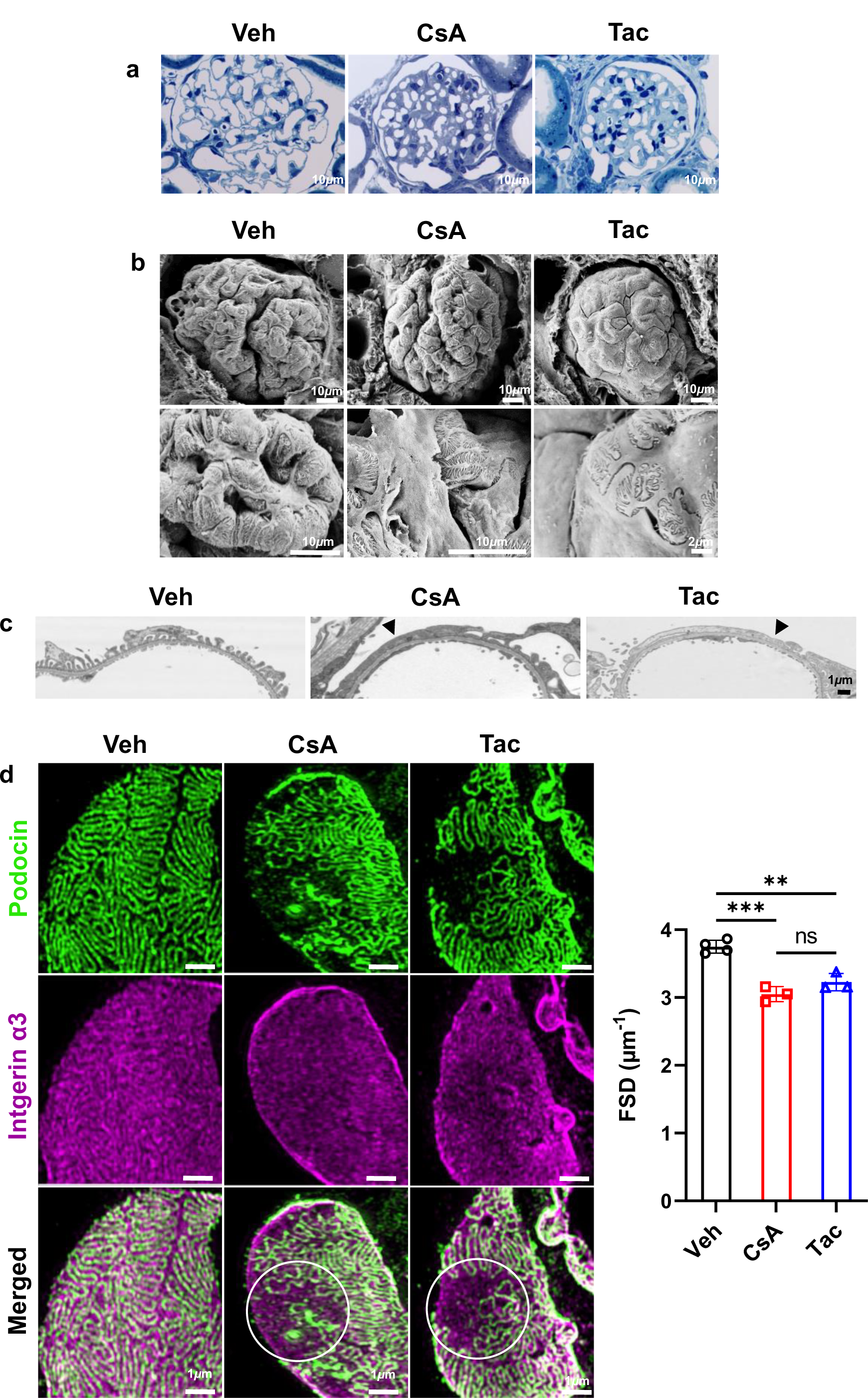
Glomerular structure. (a) Representative glomeruli, semithin sections, Richardson’s stain. Note the decreases of glomerular tuft size in the calcineurin inhibitor-treated samples. Tuft area, quantified from podocin-immunostained sections, was 7880±350 µm^2^ in vehicle (Veh) compared to 6470±600 µm^2^ in cyclosporine A (CsA) and 6820±350 µm^2^ in tacrolimus (Tac) (means ± SD; \*\**P*<0.01); difference between CsA and Tac was not significant. (b) Representative SEM of glomerular tufts as viewed from the urinary space after removal of Bowman’s capsule. Note single capillaries with flattened podocytes and retracted or fused foot processes in CsA compared to generalized effacement of podocytes with broad cell bodies and rarified foot processes in Tac. (c) Representative TEM images of the filtration barrier. Note foot process effacement in the CsA and Tac samples (arrowheads). (d) Filtration slit density (FSD) was determined by 3D-structured illumination microscopy and Podocyte Exact Morphology Measurement Procedure; double-immunostaining with anti-podocin antibody to label the filtration slit and anti-integrin α3 antibody to label the glomerular basement membrane. Flat sections of capillary loops reveal sites of effacement in CsA and Tac (white circles in the merge images) by missing slit diaphragm of portions otherwise reactive for integrin α3. Graph shows numerical evaluation; values are means ± SD; \*\**P*<0.01; \*\*\**P*<0.001. Bars indicate magnification. Statistical tests were performed using ANOVA with Tukey’s multiple comparison test.

**Figure 4.**
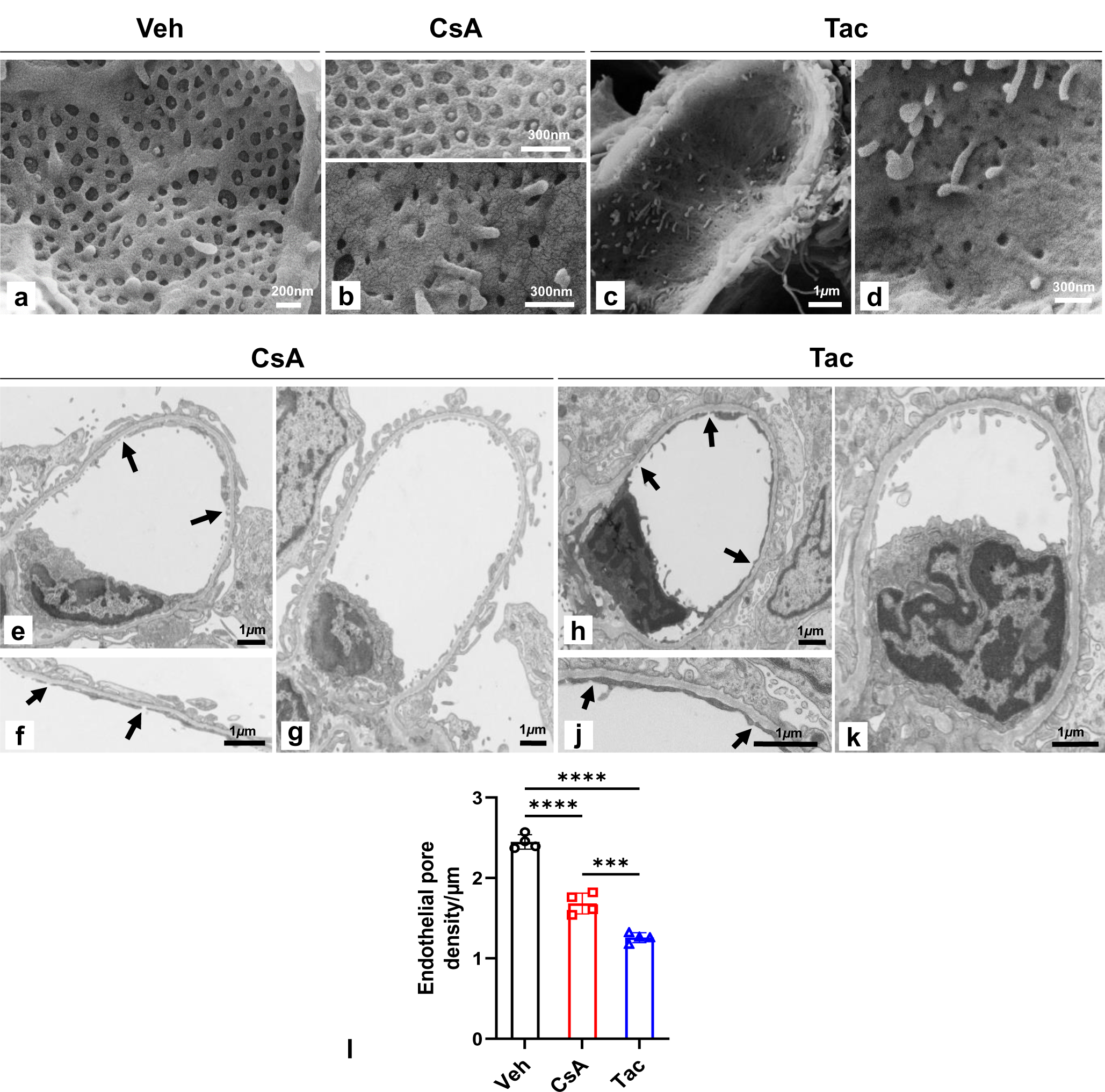
Glomerular capillaries - endothelial pore density. (a-d) Representative SEM images of glomerular capillary endothelial structure. Normal (top) or reduced pore density in cyclosporine A (CsA) (below, b). Strongly diminished pore density in tacrolimus (Tac) (overview, c; detail, d). Note also smaller pore size in the calcineurin inhibitor-treatedsamples. (e-g) TEM showing characteristic capillary endothelial structure in CsA; capillaries with (between arrows; e,f) and without obvious pore losses (g). Detail in (f) is from another capillary. Note otherwise similar endothelial morphology in (e) and (g). (h-k) In Tac, capillaries with (between arrows; h,j) and without pore losses (k) are detectable in parallel. Note striking condensation and darkening of endothelium when displaying pore losses (h). Bars indicate magnification. (l) Endothelial pore density per µm of glomerular basement membrane quantified from TEM sections; values are means ± SD; *\*\*\*P*<0.001, *\*\*\*\*P*<0.0001. Statistical tests were performed using ANOVA with Tukey’s multiple comparison test.

**Figure 5.**
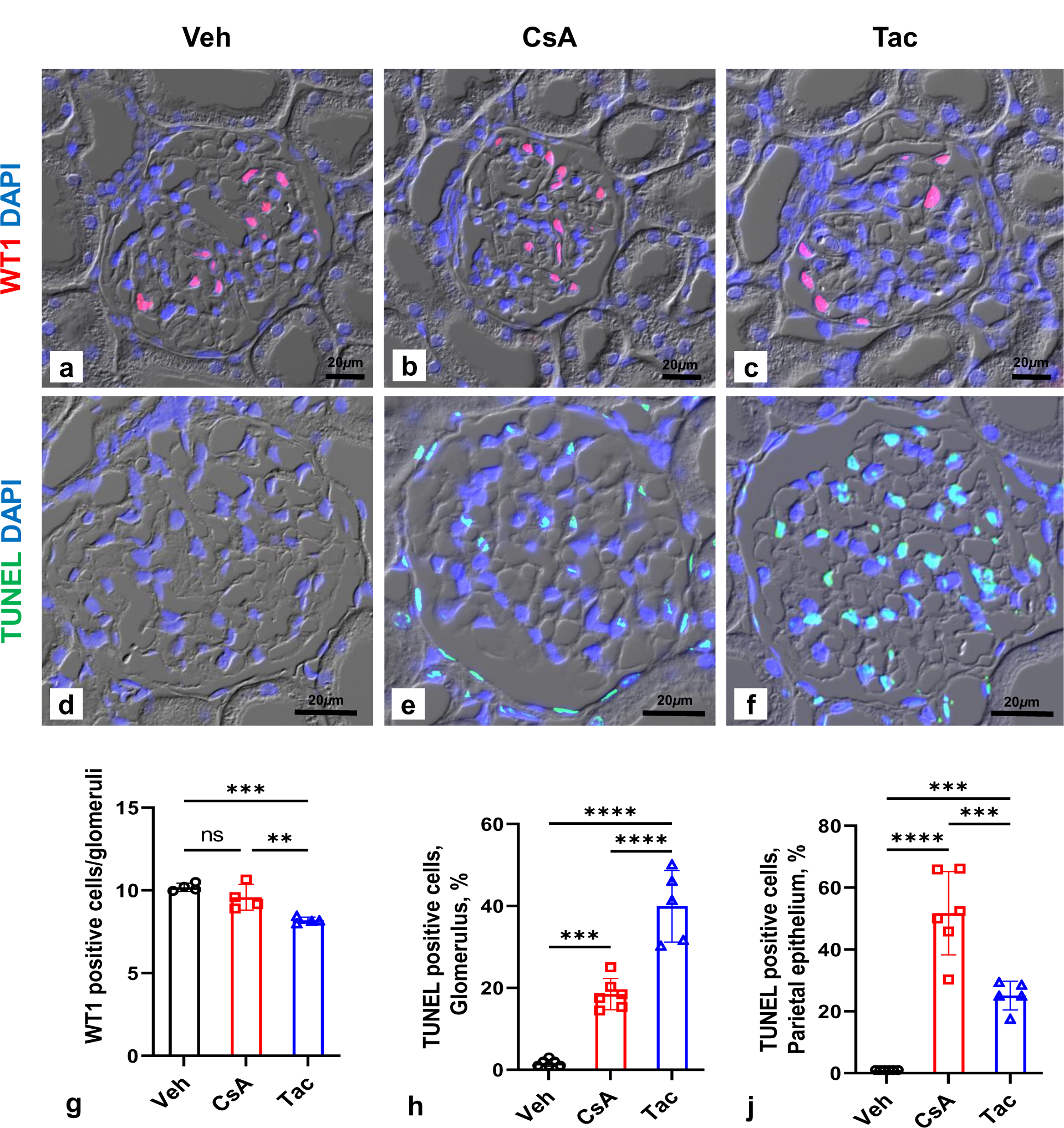
Glomerular Wilms tumor 1 protein (WT1) expression and TUNEL labeling. (a-c) Anti-WT1 immunoreactivity in podocyte nuclei. (d-f) TUNEL immunofluorescence in glomeruli. Glomerular tuft and parietal epithelium show signals in cyclosporine A (CsA) and tacrolimus (Tac). DIC optics, DAPI nuclear stain (a-f); bars indicate magnification. (g-j) Quantification of signals; values are means ± SD; \*\**P*<0.01; \*\*\**P*<0.001; \*\*\*\**P*<0.0001; ns, not significant. Statistical tests were performed using ANOVA with Tukey’s multiple comparison test.

### Proximal tubule

A variety of early changes in proximal convoluted tubule (PCT) were registered upon CNI administration. In CsA, typical Richardson’s dark-stained lysosomes were largely absent from PCT but instead, one to several large heteromorphic vacuoles as well as autofluorescent residual bodies were commonly detected per cell. The vacuoles reached nuclear size or beyond and were filled with proteinaceous content verified by PAS. In Tac, dark-stained lysosomes were less than in Veh, and heteromorphic vacuoles were rare. Lysosomal changes were quantified (Fig. 6a-b). By TEM, membrane-bound heteromorphic vacuoles, focally opening into the cytosol or anastomosing with other lysosomes, residual bodies with granular content, and late endosomes were identified (Fig. 6c). Stages of lysosomal exocytosis were frequent (Suppl. Fig. S6a-d). This was confirmed by SEM, showing the highest numbers of cast-like remainders of lysosomal exocytosis within the brush border membrane (BBM) of CsA samples (Suppl. Fig. S6e-h). The lysosomal nature of the heteromorphic vacuoles was identified by lysosomal associated membrane protein 1 (LAMP1) immunoreactivity forming a luminescent ring along each late endosomal/lysosomal perimeter in the CNI groups (Fig. 6d). Abundance of catalase as an indicator of defense against oxidative stress was elevated in CsA compared to Veh, whereas in Tac there was no difference; these changes were quantified (Suppl. Fig. S7a-d,j). Major clusters of peroxisomes were observed located in basal cell aspect (Suppl. Fig. S7e-h). TUNEL immunofluorescence showed clear maxima of nuclear signals in PCT and proximal straight tubule (PST) in CsA as opposed to few scattered signals registered in Tac (Fig. 7a-f). This was confirmed by numerical quantification (Fig. 7g-j). Further progression to necrosis affected proximal segments in both groups similarly, with higher lipid accumulations in Tac.

**Figure 6.**
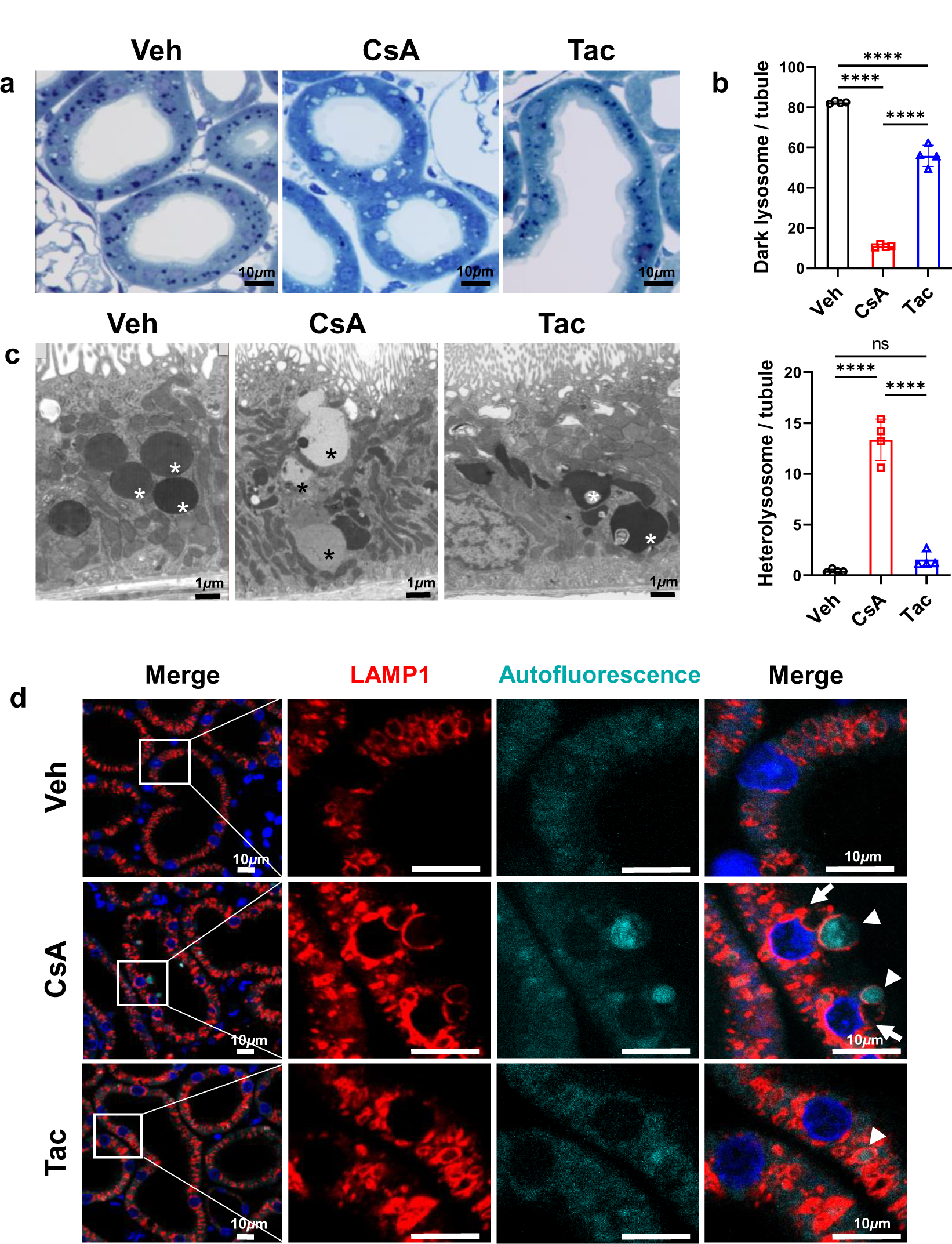
Lysosomal changes in proximal convoluted tubule. (a) Richardson‘s staining, semithin sections. Proximal convoluted tubule (S1 and S2 segments) shows well-preserved standard morphology in vehicle (Veh) with 3 to 15 dark-blue stained lysosomes per sectioned cell. Contrastingly, cyclosporine A (CsA) presents with large, clear heterolysosomes and tacrolimus (Tac) with diminished number of dark lysosomes. (b) Quantification of dark-stained lysosomes (top) and heterolysosomes (bottom); values are means ±SD; \*\*\*\**P*<0.0001; ns, not significant. Statistical tests were performed using ANOVA with Tukey’s multiple comparison test. (c) TEM; apical vesicular compartment, endosomes, fields of dense apical tubules, and autophagic vesicles with membraneous content are normally developed in Veh. Note multiple heterolysosomes (asterisks) forming chains in apico-basal orientation in CsA. In Tac, note multiform lysosomes with heterogenous inclusions in Tac (asterisks). (d) Anti-Lysosomal-associated membrane protein 1 (LAMP1) immunofluorescence of lysosomal stages; boxes in overviews show details on the right; note large heterolysosomes with LAMP1-positive perimeters irrespective of autofluorescent (arrowheads) or non-autofluorescent content (arrows) in CsA; both selected heterolysosomes are in the process of luminal exocytosis. In Tac, large heterolysosomes and autofluorescent inclusions occur only rarely (arrowhead). DAPI nuclear stain. Bars indicate magnification.

**Figure 7.**
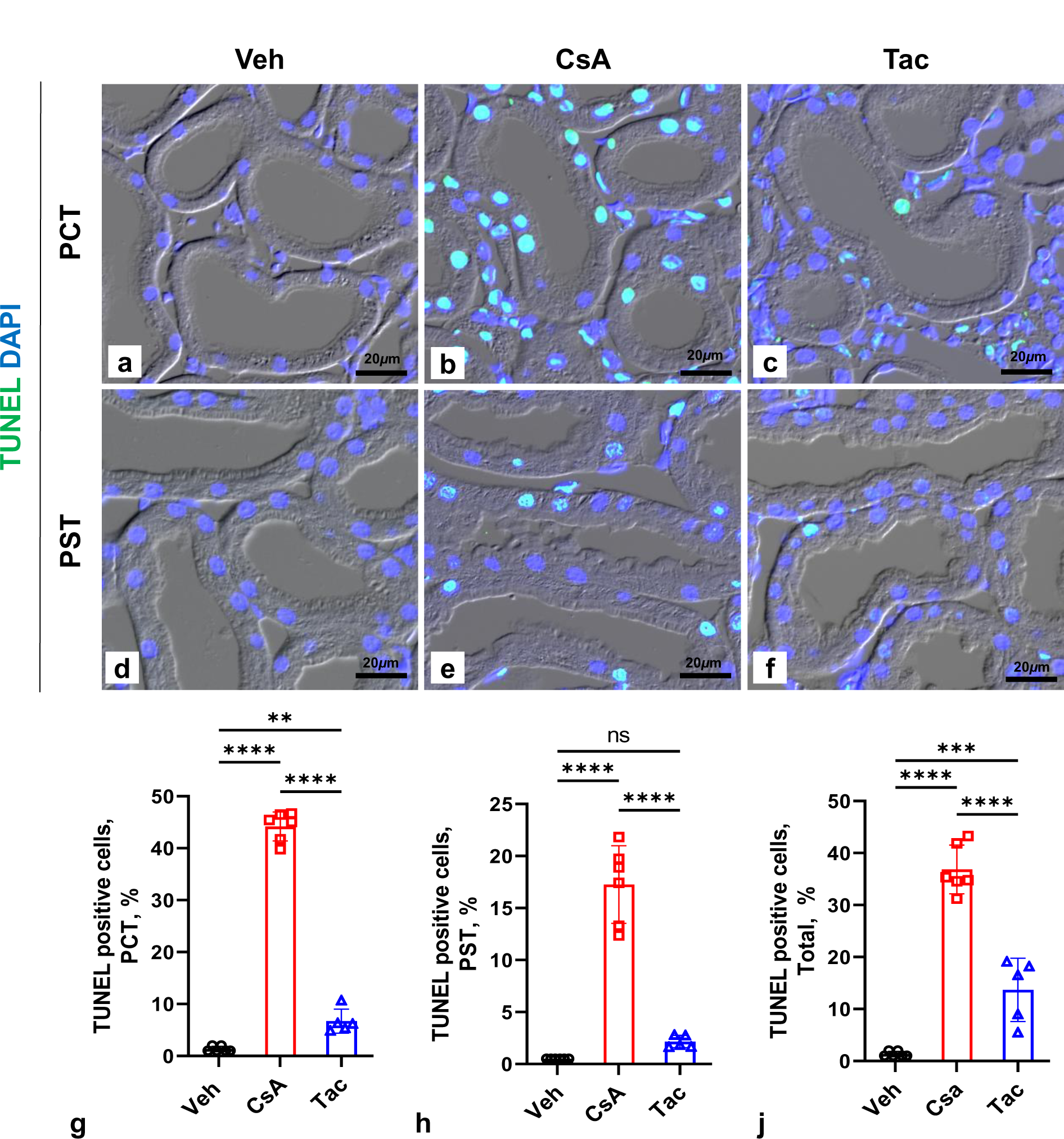
Proximal tubular TUNEL labeling. (a-f) TUNEL immunofluorescence; shows green nuclear signal in representative images. Note strong and frequent signal in cyclosporine A (CsA) (b) and weak scattered signal in tacrolimus (Tac) proximal convoluted tubule (PCT) (c). Similar results in proximal straight tubule (PST) with less signal density in CsA (E). DIC optics, DAPI nuclear stain; bars indicate magnification. (g-j) Quantitative evaluation; total kidney TUNEL signal is shown for comparison (j). Percent of total nuclei counts; values are means ± SD; \*\**P*<0.01***; \*\*\*\**P*<0.0001; ns, not significant. Statistical tests were performed using ANOVA with Tukey’s multiple comparison test.

### Changes in Cortical Interstitial Vasculature

Intact tissue areas of the CsA and Tac groups showed regular capillary CD31 signal, whereas in fibrotic foci, numerous strongly CD31-positive capillaries of a rounded, sprouting type were detected (Fig. 8a-c). Increases of CD31 abundance in CNI groups were confirmed by Western blot analysis (Fig. 8d). Changes in endothelial fenestration were quantified by TEM; CNI diminished the pore density significantly with decreases stronger in Tac than in CsA (Fig. 8e-h).

**Figure 8.**
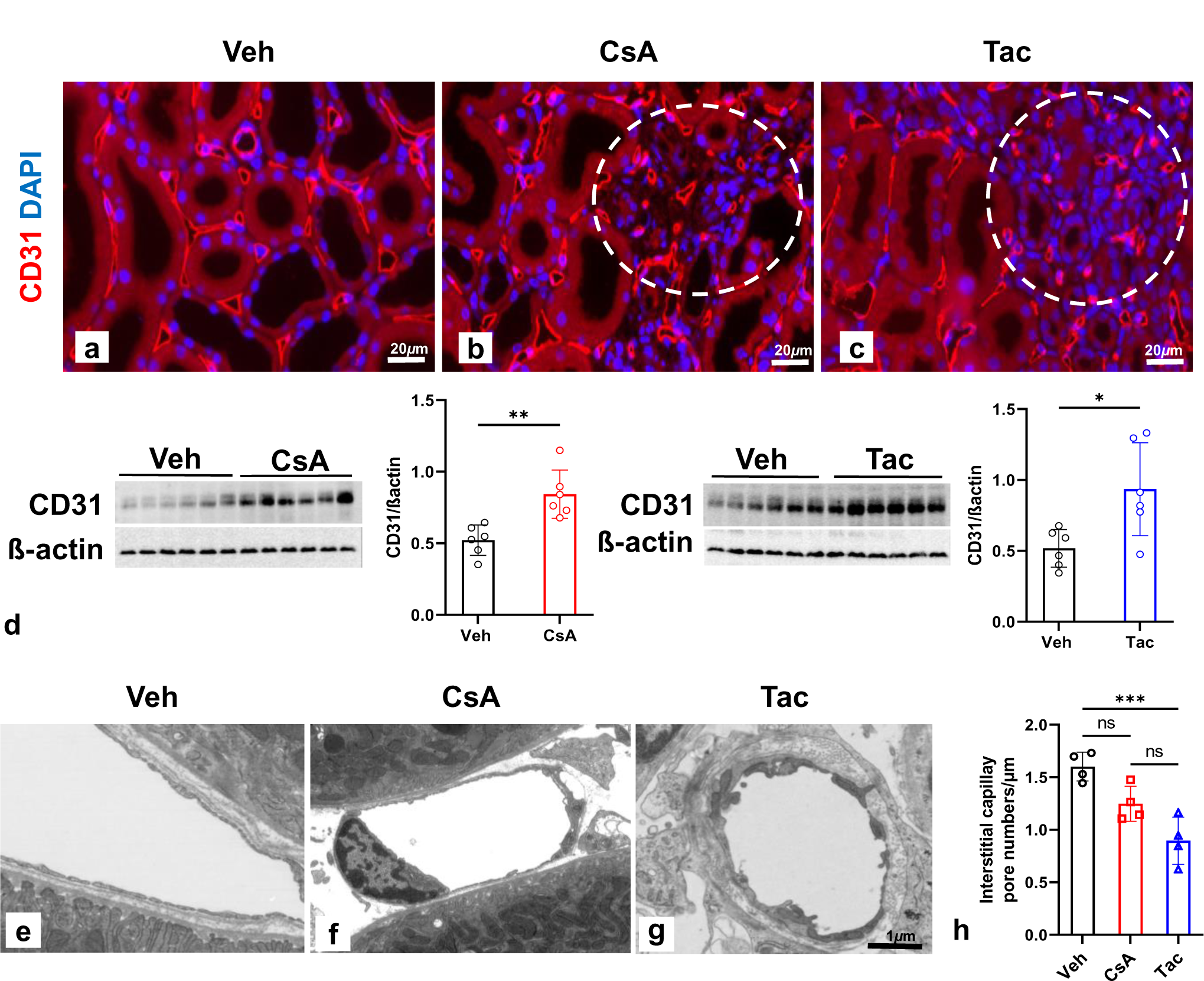
Changes in cortical interstitial vasculature. (a-c) Anti-CD31 immunofluorescence localized to peritubular capillaries. Note strongly positive capillaries of a rounded, sprouting type in fibrotic foci (encircled areas in b,c); DAPI nuclear blue staining. (d) Western blot CD31 signals (110 kDa) from kidney extracts of cyclosporine A (CsA), tacrolimus (Tac), and their respective vehicle (Veh) groups (n=6, respectively); ß-actin (42 kDa) serves as loading control. Densitometric evaluations on the right; data are means ± SD; \**P*<0.05; \*\**P*<0.01. (e-g) TEM of cortical interstitial capillaries in fibrotic foci. Note sprouting type capillaries with absent fenestrae in CsA and Tac, and endothelial swelling or ridge formation particularly in Tac (g). (h) Endothelial pore density per µm of capillary basement membrane quantified from TEM sections; values are means ± SD; *** *P*<0.01; ns, not significant. Bars indicate magnification. Statistical tests were performed using unpaired Student’s *t* test (d) or ANOVA with Tukey’s multiple comparison test (h).

### Multiomics Analyses

RNA-seq transcriptomic, global proteomic, and phosphoproteomic analyses served to obtain mechanistic perspectives underlying CNI-induced pathology. RNA-seq revealed 1003 differentially expressed genes (DEG; 507 up- and 496 downregulated) in CsA and 323 (105 up- and 218 downregulated) in Tac. Few products were jointly altered, among them increased renin and decreased calbindin (adjusted *P*<0.1; see Venn diagram in Suppl. Fig. S8a). Pathway enrichment analysis of CNI-induced DEGs by Enrichr indicated that among others, *vasculature development*, *glomerular visceral epithelial cell differentiation, actin cytoskeleton organization*, *eukaryotic translation termination*, and *apoptotic process* were upregulated, whereas *proximal tubule transport* and likely associated metabolism were downregulated, supporting CsA-typical damages. In Tac, *renin-angiotensin system*, *biological oxidations*, *oxidative stress-induced senescence*, *blood vessel endothelial cell proliferation involved in sprouting angiogenesis*, and *vascular wound healing* were up-, and *vasculature development*, *positive regulation of endothelial cell proliferation*, *mTORC1 signaling*, *AMPK signaling pathway*, and *actin filament organization* downregulated, which agreed with Tac-typical structural alterations (Fig. 9). Both proteomic and phosphoproteomic data were consistent with these pathways and further selectively pointed to markedly upregulated *unfolded protein response* pathway in CsA but not in Tac; the respective products were verified by WB (Suppl. Fig. S9-11). Among single, differentially regulated products, PDIA5 and phosphorylated PERK (*Eif2ak3*), related with UPR, as well as HMGB1 and CTSD, both regulatory in autophagy, were upregulated in CsA. In Tac, *Prkce, Prkd1, Ptger4*, and *Cav1* transcripts, CAV1 and DDAH2 proteins, and phosphorylated PRKD1, all involved in endothelial function and vasculature changes, were downregulated. Likewise, SYNPO transcript and protein, PALLD transcript and phosphorylation, PODXL protein, and CD2AP phosphorylation, all relevant in podocyte biology, were decreased. We furthermore used the published list of 193 proximal tubule-specific genes identified in rat kidney to compare with the present changes^28^. Here, 100 differentially regulated genes (20 up- and 80 downregulated) were altered in CsA, but only 33 genes (25 up- and 8 downregulated) in Tac, confirming the more substantial changes in CsA compared to Tac (Suppl. Fig. S12).

**Figure 9.**
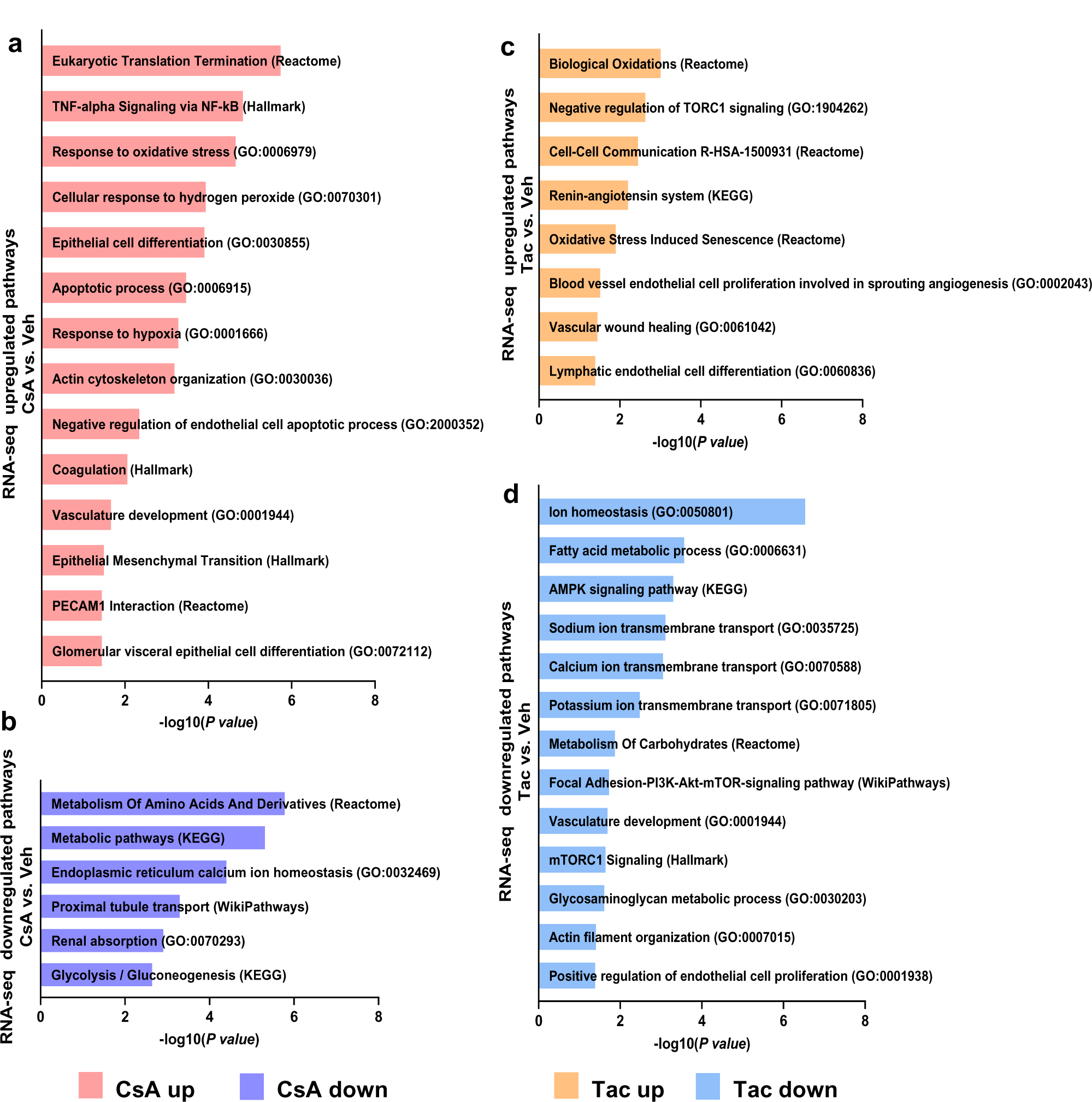
Pathway enrichment analysis of differentially expressed genes affected by calcineurin inhibitor (CNI) treatments. Treatment with cyclosporine A (CsA) and tacrolimus (Tac) leads to different molecular pathway responses in chronic CNI nephrotoxicity. Significantly enriched pathways in genes upregulated (a) and downregulated (b) in CsA versus vehicles (*P* < 0.05). Significantly enriched pathways in genes upregulated (c) and downregulated (d) in Tac versus vehicles (*P* < 0.05). Term lists used in this analysis were GO_Biological_Processes, WikiPathways, Reactome, KEGG, and Hallmark to determine enriched processes and pathways from differentially expressed genes (Enrichr webtool).

## Discussion

The presented study provides potential mechanisms of CNI nephropathy under early chronic conditions, focussing distinctive pathogenesis for CsA vs. Tac. High-resolution histopathology, candidate RNA and protein assessments as well as comprehensive multiomics attributed renal microenvironmental remodeling to novel, specific protein signatures. Responses to treatments were distinctive, associating CsA with a tubular and Tac with a microvascular focus. We share potential biomarkers related to both drugs. Pathological changes point to distinct onset changes in arteriolar wall changes of the two groups.

Greater RAAS stimulation in Tac vs. CsA may contribute to vascular injury^13, 29^. Disproportionate impairment of vasodilatory prostaglandins in Tac agrees with downregulated of Ep4 receptor (*Ptger4*) and related PGE2 action, implying less eNOS function^30, 31^. *Prkce* mRNA, selectively downregulated in Tac, may suggest impaired VEGF signaling, since its knockdown in bovine endothelial cells abrogate VEGF-stimulated AKT phosporylation, decreases action of its cognate receptor VEGFR2, and impairs VEGF-stimulated NOS activity^32^. Likewise, CAV1 mRNA & protein were downregulated in Tac but not in CsA, supporting related NO deficiency downstream of impaired VEGF signaling^33^. Selective upregulation of DDAH2, which degrades endogenous NOS inhibitors such as asymmetric dimethylarginine (ADMA) and monomethyl arginine (L-NMMA), indicate potential compensation to preserve NO availability in Tac^34^. Increased leukocyte adhesion, more in Tac than in CsA, pointed to affection of the endothelial surface and agreed with focal incidence of CD45-positive cell accumulations^35^. This difference is in line with CsA’s beneficial effects on adhesion molecules *in vitro*^36^. Stimulated endothelial CD31 in both conditions suggests a cytoprotective response to altered shear stress^37, 38^ and/or adjustment for leukocyte transmigration^39^.

In the glomeruli, Bowman’s capsules show fibrotic alterations in early and advanced foci by CsA and Tac. Pronounced CD44 expression, as found in the activated parietal epithelium, is a useful marker for early CNI-NTX in human grafts^40^. In conjunction with fibrous podocyte synechiae and CD45-positive inflammatory cell accumulations these features suggested early stages of sclerotic lesions^41, 42^. Tuft retraction, likely resulting from diminished microvascular perfusion^43^, and decreased FSD occurred to similar extent in CsA and Tac. Clearly, however, endothelial fenestration is affected more by Tac than CsA in glomerular as well as peritubular capillaries and renal veins. VEGF signaling involvement appears possible due to its effects on pore density in conjunction with TNFα signaling in mice^44^. A human microvascular CNI-NTX model further underscores VEGF’s potential role. That model displays reduced fenestration and VEGF signaling along with stimulated *Adamts1* and cytoplasmic deteriorations^45^. Accordingly, we find exacerbated endothelial cytoplasm condensation for Tac. *Adamts1 and* transcription factor *Sox17* are likewise stimulated. *Sox17* status differed between our groups and may provide compensatory endothelial repair in CsA, yet failing to do so in Tac^46^. With caveolin-1 (*Cav1*) substantially diminished at the mRNA and protein level in Tac, our multiomics data further link reduced angiogenesis to pore formation^47, 48^. This is further supported by protein kinase D1 (PRKD1) mRNA and phosporylation being selectively downregulated in Tac. PRKD1 also controls VEGFR2 by disinhibiting transciption factor AP2^49^. Extensive foot process effacement in Tac, more than in CsA, suggests impaired filtration barrier and hydraulic conductivity, since capillary segments with advanced pore rarefaction show parallel podocyte effacement. This agrees with decreased WT1 and increased TUNEL signals in podocytes, both exacerbated in Tac. Diminution of WT1, an established marker of podocytes, is associated with barrier function loss^50^ and early changes towards glomerulosclerosis^51^, while enhanced TUNEL signals relate to apoptosis or repair^52^. Key podocyte genes such as palladin, CD2AP, synaptopodin, nephronectin, and podocalyxin, all intrinsic structural components of podocytes, were selectively downregulated in Tac which also underscores higher podocyte damage compared to CsA, and concomitant impairment in shape and motility of their foot processes^53, 54^. Associated decrease in phosphorylation of CD2AP, a stabilizing protein of the slit diaphragm, at Ser 404 may reflect further loss of function^55^. Although in their longitudinal histological cohort study, Nankivell and colleagues reported equal glomerular damage scores in CsA and Tac samples^3^, our results suggest glomerular vulnerability to Tac at an early stage.

Tubular damage in CNI nephropathy has been extensively discussed in the past, yet it has remained uncertain whether whether CNI damage occurs by its toxicity to epithelia, downstream effects of affected vasculature, or glomerular function prevail at its origin. The proximal tubule is considered to be the principal nephron segment affected by CsA and Tac, with no distinction^56, 57^. In contrast, the presented data suggests substantial differences in proximal tubular damage. The major finding is massive accumulation of heterolysosomes, identifed by their LAMP1-positive boundaries, and their exocytotic luminal extrusion predominantly in the CsA group, whereas changes in Tac were much less lysosomal-based. Our multiomics data confirm milder tubulotoxicity by Tac, in harmony with Tac’s preferential use in organ transplantation^1, 3^. PERK is a key regulator and component of the UPR that markedly increased together with the UPR transducer IRE1 and downstream spliced XBP1. These canonical cascades likely prompted cleaved caspase-3 and related apoptosis shown here for CsA, but not for Tac^18, 58^. Already before, we and others reported impaired proteostasis and UPR leading to proapoptotic ER stress upon CsA, but not for Tac^17, 20^. Several lines point to the localization of these particular pathways in the proximal tubule. Among them are the TUNEL signal and related apoptotic cell features^20, 58^. With the UPR response also driving regulation of TFEB, an acknowledged master regulator of autophagy and lysosomal biogenesis^59^, CsA may block autophagic flux, thus causing the selective lysosomal disorder that was not observed in Tac. Notably, mTOR inhibitors, commonly employed in combination with CNI, may have a beneficial effect herein through their activation of TFEB^22, 59, 60^. Stimulated *Arl8b* expression standing for enhanced lysosomal exocytosis, which we observed in the proximal tubule of the CsA group, supported the lysosome-based defect^61^. Upregulated HMGB1 and CTSD indicate bioenergetic maladaption and mitochondrial dysfunction along with enhanced autophagy in CsA^57, 62, 63^. Such disorders may arise with oxidative stress, since CsA stimulated catalase with activated proximal tubular peroxisomes^64^.

It is therefore reasonable to conclude that CsA-related cytotoxicity, among other sites, preferentially affects these proximal tubule tissue components. Comparison of our DEG signatures with published library data referring to the proximal tubule support this interpretation (Suppl. Fig. S12). The presented changes typically begin at the urinary pole and affected the entire length of PCT, but had little structural manifestation in PST. This is unlike early findings in the rat suggesting the PST as the primary target for CsA toxicity.^56^ Higher CsA dosage, combination with low salt diet, and shorter treatment may account for these differences. Apoptosis in PCT was markedly higher in CsA than in Tac, but also included PST, suggesting damage potential in the S3 segments as well. Further advanced damage features indicated progression to similar loss of proximal tubular differentiation in either group.

In summary, we show distinctive pathogenetic patterns arising in renal compartments under CsA vs. Tac medications. Besides common effects like fibrosis and atrophy, landmark differences were the predominant affections of the filtration barrier in Tac and of the PCT in CsA. Protein signatures from selective probing and multiomics analysis point at the mechanisms involved. For Tac, these include perturbed VEGF/VEGFR2 signaling. Cytotoxic UPR signalling and altered autophagic flux is more prominent for CsA. Products identified from multiomics provide differential pathognomonic biomarkers, as ranked and assigned in the *graphical abstract*. Biomarker multiplexed detection may allow early diagnosis and prognosis of allograft failure. In the future, CNI selection may increasingly depend on individual patient risk factors for damage of the filtration barrier versus the proximal convoluted tubule.

## Data Statement

EM large-scale digitization data are available under www.nanotomy.org repository. The RNA-seq data sets are available from the Gene Expression Omnibus repository under the following accession numbers: GSE225215. Proteomic and phosphoproteomic data sets are available via ProteomeXchange with identifier PXD038841, PXD038546, respectively.

## Acknowledgements

This work was financially supported by Deutsche Forschungsgemeinschaft BA700/22-2, MU2924/2-2, and SFB 1365-C04/-S01. HD was supported by a doctoral fellowship from Ministry of National Education, Turkey. We thank Hermann-Josef Gröne, Wilhelm Kriz, David Ellison, and Richard Warth for constructive advice, Anette Drobbe for secretarial help, Kerstin Riskowsky, Katja Dörfel, Ariane Anger for expert technical help, Sara Timm, Petra Schrade, and John Horn (Core Facility for Electron Microscopy) for microscopical help, Junda Hu, Erdmann Seeliger, Nicole and Tim Endlich (NIPOKA, Greifswald, Germany) for methodological help in 3D-SIM, and Lukasz Szyrwiel, Philipp Mertins, and Nils Blüthgen for bioinformatic data analysis.

## Author Contributions

SB and HD conceived and designed research; SB, HD, KM, and SP supervised animal experiments and tissue processing; HD, SB, SP, DEY, CD, and KM performed and analyzed histological, physiological, and biochemical experiments; HD, IAE, MM, CH, and SB designed and analyzed multiomic data; SB and HD designed figures; SB and HD drafted manuscript; SB, HD, CH, and PBP approved final version of manuscript.

## SUPPLEMENTARY MATERIAL

Supplementary File (PDF)

### 1. Supplementary Methods

### 2. Supplementary Figures

**Supplementary Figure S1: Smooth muscle actin (α-SMA) and leukocyte common antigen (CD45) staining.** (a-g) Anti-α-SMA immunoperoxidase staining in vehicle (Veh) (a,e) shows vascular wall signals. In cyclosporine A (CsA) and tacrolimus (Tac), interstitial signals are enhanced in foci around tubules and glomeruli, and in outer stripe focal areas. Signal is pronounced near atrophic or necrotic tubules and adjacent glomeruli, and pericapsular expression is variably present with stronger signals encountered in the CsA than in the Tac group (b, c, f, g). Bar graphs indicate average size of α-SMA-stained cortical foci (percent of unit sectional area; d); values are means ± SD; \*\*\**P*<0.001; ns, not significant. (h-k) CD45 immunofluorescence staining shows scattered individual interstitial cells in the cortex of Veh (h) and enhanced interstitial signals in fibrotic foci of CsA and Tac samples (j, k). DIC optics; bars indicate magnification. Statistical tests were performed using ANOVA with Tukey’s multiple comparison test (d).

**Supplementary Figure S2: Renal arterial and arteriolar wall structure.** (a-c) Interlobular arterial walls are compared in representative views; in a vehicle (Veh) sample, endothelium, media, and adventitia present normal structure (a). In cyclosporine A (CsA), occasional vesicular or hyaline inclusions, here in subintimal location (asterisk in b) are shown. In tacrolimus (Tac), thinning of media wall cells, augmented matrix deposition, and thickened adventitia with substantial fibrillar collagen I layers (open circle) are characteristic (c). (d-g) Afferent arterioles, Periodic acid-Schiff (PAS); note the distinct staining intensities; luminal occlusion was generally not observed. (d-f). Wall-to-lumen ratios (g); values are means ± SD; ** *P*<0.01; ns, not significant. (h-k) Wall structure and endothelial CD31 immunoreactivity; note stronger signal in CNI groups. (l-n) Afferent arteriolar wall structure by TEM. DAPI blue nuclear staining and DIC optics (h-k); bars indicate magnification. Statistical tests were performed using ANOVA with Tukey’s multiple comparison test (g).

**Supplementary Figure S3: Juxtaglomerular apparatus – renin. (**a-d) Anti-renin immunoperoxidase staining shows normal distribution of renin signal in the preglomerular afferent arteriolar portion (a), highly increased signal in cyclosporine A (CsA) (b), and even higher increase in tacrolimus (Tac) (c) in representative views. In Tac, narrowing of the preglomerular afferent arteriolar portion of retracted-appearing glomeruli occurs (d); here, granular cells may form layers. (e) Bar graphs show quantitative expression of renin (immunoreactive area as percent of unit sectional area); values are means ±SD; * *P*<0.05, **** *P*<0.0001. (f-h) TEM shows high glycogen deposits in juxtaglomerular granular cells next to the renin granules in Tac (g) compared to CsA (f), and necrotic inclusions may be observed in granular cell fields in Tac (h). DIC optics, hematoxylin stain (a-d); bars indicate magnification. Statistical tests were performed using ANOVA with Tukey’s multiple comparison test (e).

**Supplementary Figure S4: Renal venous endothelia.** (a-e) Representative SEM views of venous endothelia (arcuate or interlobular veins); typical distribution of fenestrated pore fields in a vehicle (Veh) sample (a); similar structure is seen in cyclosporine A (CsA) next to a pore-free field (b), whereas in tacrolimus (Tac), normal pore fields (c) or larger areas of absent fenestration may be encountered (d); adherent leukocytes are frequent (e). (f-h) CD45 immunoreactivity shows scattered perivascular leukocytes in Veh (f), few adherent leukocytes in CsA (g), and numerous adherent leukocytes in Tac (h). DAPI blue nuclear staining and DIC optics (f-h). Bars indicate magnification.

**Supplementary Figure S5: Changes in Bowman’s capsule.** (a-f) TEM representative images show regular structure of Bowman‘s capsule in vehicle (Veh) with flat parietal epithelium and thin capsular basement membrane (a). In cyclosporine A (CsA), both parietal epithelium and capsular basement membrane are markedly thickened, ranging from 0.2 up to 4 µm in cell height, with extensive granular and hyaline matrix formation (b), and synechiae with multiple contact points are formed (c); podocytes may form synechiae even in the urinary pole area where they may display tip lesions (d,e). In tacrolimus (Tac) a similar synechia between podocyte and parietal epithelium is shown; note typical rER inclusion body within podocyte (asterisk) (f). (g) Anti-CD44 immunofluorescence staining shows occasional mild signal in Bowman‘s capsule in a Veh sample, whereas in CsA and in Tac activated parietal epithelium presents with high signal intensity along almost the entire perimeter. (h) Anti-CD45 staining shows few interstitial signals in Veh, but strong signal associated with the capsule in CsA and, more so, in Tac. DAPI blue nuclear staining and DIC optics; bars indicate magnification.

**Supplementary Figure S6: Ultrastructure of lysosomal canges in proximal tubule.** (a-d) TEM reveals that in cyclosporine A (CsA), heterolysosomes reveal partial communication with the cytosol via interrupted membrane portions (arrows; a), anastomose with residual bodies (arrow) and late endosomes (asterisk; b), are filled with heterogenous debris, reach the luminal cell membrane (c), and are frequently encountered in exocytotic transition (d). (e-h) Representative SEM views show the frequency of lysosomal exocytosis in proximal convoluted tubule by the number of transitory states, leaving pits in the brush border. Vehicle (Veh) defines is baseline activity (e), whereas in CsA exocytosis is substantially enhanced (f); in tacrolimus (Tac) there is intermediate activity (g); detailed view of pit formations (arrows; h). Bars indicate magnification.

**Supplementary Figure S7: Catalase abundance and peroxisome structure in the proximal tubule.** (a-d) Immunofluorescence staining with anti-catalase antibody. There is abundant staining in proximal convoluted tubule of Vehicle (Veh) (a), cyclosporine A (CsA) (b), and tacrolimus (Tac) samples (c). Inserts reveal punctuate nature of the signal; DAPI blue nuclear staining. Note intensified signal in CsA. Bar diagram shows enhanced fluorescence signal quantified per unit sectional area from catalase-immunostained sections; values are means ± SD; * *P*<0.05; ns, not significant. (e-h) TEM images showing regular distribution of peroxisomes at the basal cell pole (e) and regular peroxisome morphology with adjacent cisternae of rough and smooth endoplasmic reticulum at higher resolution in Veh (f). Clusters of peroxisomes are visualized in CsA (g) and Tac (h); their extent may vary with treatment and cell affection. (j) Western blots show immunoreactive catalase (60 kDa) from kidney extracts of CsA, Tac, and their respective vehicle groups (n=6, respectively); ß-actin (42 kDa) serves as loading control. Densitometric evaluation is presented on the right side of the respective blots. Data are means ± SD; *** *P*<0.001, ns, not significant. Statistical tests were performed using ANOVA with Tukey’s multiple comparison test (d) or unpaired Student’s *t* test (j). Cat, catalase.

**Supplementary Figure S8: Differentially expressed genes, proteins, and phosphoproteins.** (a-c) Venn diagram showing the number of differentially expressed genes, proteins, and phosphoproteins (RNA-seq and proteomics, adjusted *P*< 0.1; phosphoproteomics, FDR<5; CNI vs. Veh). Jointly regulated products arelisted in box. “up”, upregulated; “down”, downregulated.

**Supplementary Figure S9: Pathway enrichment analysis of differentially expressed proteins (DEPs) affected by CNI treatments.** Significantly enriched pathways in proteins up- (a) and downregulated (b) in cyclosporine A (CsA) versus vehicle (Veh) (*P* < 0.05). Significantly enriched pathways in proteins up- (c) and downregulated (d) in tacrolimus (Tac) versus vehicle (Veh) (*P* < 0.05). Term lists used in this analysis were GO_Biological_Processes, WikiPathways, Reactome, KEGG, and Hallmark to determine enriched processes and pathways from DEPs (Enrichr webtool).

**Supplementary Figure S10: Pathway enrichment analysis of differentially expressed phosphoproteins (DEPPs) affected by calcineurin inhibitor treatments.** Significantly enriched pathways in phosphoproteins up- (a) and downregulated (b) in cyclosporine A (CsA) versus vehicle (Veh) (*P* < 0.05). Significantly enriched pathways in phosphoproteins upregulated (c) and downregulated (d) in tacrolimus (Tac) versus vehicle (Veh) (*P* < 0.05). Term lists used in this analysis were GO_Biological_Processes, WikiPathways, Reactome, KEGG, and Hallmark to determine enriched processes and pathways from DEPPs (Enrichr webtool).

**Supplementary Figure S11: Western blot verification of unfolded protein response parameters.** Phospho-PKR-like ER kinase (p-PERK,Thr980,150 kDa), phospho-Inositol requiring-1α (p-IRE1α, Ser724, 110 kDa), X-box binding protein 1 (sXBP1) (55 kDa), cleaved caspase 3 (c-casp3) (17 kDa), and ß-actin (42 kDa), rat kidney. Below, corresponding densitometric evaluations expressed as –fold changes relative to vehicle (Veh) levels. Data are means ± SD; *n*=6 per group. \*\*\**P* < 0.001, \*\*\*\**P* < 0.0001; ns, not significant. Statistical tests were performed using unpaired Student’s t test.

**Supplementary Figure S12: Differentially expressed genes with endogenous expression in proximal tubule.** Bar graph on a published RNA-seq data set of segmental profiles of the adult rat nephron ^s^^9^ showing 193 genes endogenously expressed in proximal tubule; 100 of these are differentially expressed in the cycslosporine A (CsA) group but only 33 in tacrolimus (Tac) (*P* value < 0.05). Here, a tenfold higher number of transcriptionally downregulated genes in CsA compared to Tac should correspond to significant loss of function in this nephron segment. They indicate a tendency towards fibrosis and chronic kidney disease in CsA, such as upregulated *Sglt2* pointing to profibrotic changes of the S1 and S2 segments ^s10^, as well as downregulated *Hao2* standing for decreased fatty acid metabolism ^s11^, *Qdpr* for TGFß1, NOX1 and NOX4 stimulation ^s12^, *Tmigd1* for impaired protection from oxidative cell injury ^s13^, *C1qtnf3* for TGFß1 and interstitial fibrosis ^s14^, and *Kmo* for epithelial injury and allograft rejection ^s15^. By contrast, in Tac stimulated *Cyp2e1* indicated activated resistance against epithelial decay which in this case may be reactive upon upstream glomerular injury ^s16^. Veh, vehicle.

### 3. Supplementary Tables

Supplementary Table S1: List of antibodies used for immunohistochemistry and Western blot.

### 4. Supplementary References

