## Supplementary material for "Immunosuppression with Cyclosporine versus Tacrolimus shows distinctive nephrotoxicity profiles within renal compartments"

**Supplementary Table of Content**

**1. Supplementary Methods.**

**2. Supplementary Figures.**

Supplementary Figure S1: Smooth muscle actin (α-SMA) and leukocyte common antigen (CD45) staining.

Supplementary Figure S2: Renal arterial and arteriolar wall structure

Supplementary Figure S3: Juxtaglomerular apparatus – renin

Supplementary Figure S4: Renal venous endothelia.

Supplementary Figure S5: Changes in Bowman's capsule.

Supplementary Figure S6: Ultrastructure of lysosomal canges in proximal tubule.

Supplementary Figure S7: Catalase abundance and peroxisome structure in the proximal tubule.

Supplementary Figure S8: Differentially expressed genes, proteins, and phosphoproteins.

Supplementary Figure S9: Pathway enrichment analysis of differentially expressed proteins (DEPs) affected by CNI treatments.

Supplementary Figure S10: Pathway enrichment analysis of differentially expressed phosphoproteins (DEPPs) affected by calcineurin inhibitor treatments.

Supplementary Figure S11: Western blot verification of unfolded protein response parameters.

Supplementary Figure S12: Differentially expressed genes with endogenous expression in proximal tubule.

**3. Supplementary Tables.**

Supplementary Table S1: List of antibodies used for immunohistochemistry and Western blot.

**4. Supplementary References.**

### Supplementary Methods

#### Animals

Animal experiments were approved by the German Animal Welfare Regulation Authorities for the protection of animals used for scientific purposes (Berlin Senate; G0148/18). Adult (10 to 12 week-old) male Wistar rats were divided into groups receiving cyclosporine A (CsA, Sandimmun, Novartis; target plasma trough level 3 µg/ml) and the respective vehicle (Veh, saline), or tacrolimus (Tac, FK506, Selleckchem; target plasma trough level 3.5 ng/ml) and the respective vehicle (25% DMSO/75% PEG500, Sigma) via subcutaneously implanted osmotic minipumps (Alzet, 2ML4) for 28 days. For pump filling, 100 mg CsA and 7 mg Tac/ml of vehicle were used. For minipump implantation, rats were anesthetized by isoflurane inhalation. An incision of the neck skin was performed and the subcutaneous tissue dilated to create a pocket for the pump. The filled pump was then inserted into the pocket and the wound closed with metal clips. On the second last day of each experiment, rats were placed in metabolic cages for 24 h with water and chow ad libitum to collect urine. At the end of the experiments, rats were anesthetized with ketamin/xylavet (90/10 mg/kg BW; CP-Pharma) to obtain blood samples. Urine and blood samples were analyzed by a commercial laboratory (IMD Labor). For biochemical evaluation, one kidney was clamped and removed for biochemical analysis before perfusion fixation, or both kidneys were removed without fixation.

#### Blood and urine analysis

Blood was taken from the inferior vena cava using heparinized syringes, decanted into Eppendorf tubes, left for 30 min at room temperature (RT) for clotting, and centrifuged at 2.000xg for 10 min at 4 °C to obtain serum with the supernatant. Creatinine was measured and its clearance calculated using the formula, CrCl (ml/min) = (urine creatinine [mg/dl] × urine flow [ml/min] / serum creatinine [mg/dl]). Fractional excretion of sodium (FeNa) was calculated by the formula, FeNa (%) = (urinary sodium [mg/dl] × serum creatinine [mg/dl]) / (serum sodium [mg/dl] × urinary creatinine [mg/dl]).

#### Perfusion fixation and tissue processing

Kidneys were perfused via the abdominal aorta, first with 3% hydroxyethyl starch in 0.1 M Na-cacodylate (Caco) for 20 to 30 sec, then with 3% paraformaldehyde/3% hydroxyethyl starch in Caco for 5 min. For paraffin embedding, tissue was post-fixed in the same fixation solution overnight at 4 °C, then transferred to Caco supplemented with 300 mOsm sucrose and 0.02% NaN3 until embedding. For cryostat sectioning, tissue was transferred directly after perfusion to 800 mOsm sucrose in Caco at 4 °C overnight, snap-frozen in 2-methyl butane cooled with liquid nitrogen, and stored at -80 °C. For histology and immunohistochemistry (IHC), tissues were dehydrated and paraffin-embedded. For electron microscopy, tissues were post-fixed overnight at room temperature in 1.5% glutaraldehyde/1.5% paraformaldehyde containing 0.05% picric acid in Caco, then in 1% osmium tetroxide/0.8% potassium hexacyanoferrate in Caco for 1.5 h at RT for transmission electron microscopy (TEM) or in 1% aqueous osmium tetroxide for scanning electron microscopy (SEM). Tissues were then dehydrated and embedded in epoxy resin for semithin sectioning and light microscopy (LM) or ultrathin sectioning and TEM analysis using standard methodology. For SEM, samples were high pressure-critical point-dried and sputter-coated.

**Histology and immunostaining**

For paraffin histology and IHC, 4 μm-thick sections were cut, mounted on glass slides, and deparaffinized. Histology was done using standard PAS, Masson's Trichrome, or Sirius Red staining protocols. For IHC, heat-induced epitope retrieval was generally performed for 6 min by cooking slides in citrate buffer (10 mM sodium citrate, pH 6.0) using a pressure cooker. Sections were blocked with 5% BSA (Serva) in TBS for 30 min at RT. Samples were then incubated overnight at 4 °C with primary antibody dissolved in 1% BSA/TBS or blocking medium (antibodies listed in Suppl. Table S1). After washing in TBS, fluorescently labeled secondary antibodies (Suppl. Table S1) were dissolved in 1% BSA/TBS or blocking buffer and incubated for 1 h at RT. Sections were then mounted in PBS-glycerol (1:9). Nuclei were stained with DAPI (Sigma). For immunoperoxidase staining, sections were incubated in methanol containing 0.3%

hydrogen peroxide ($H_2O_2$) for 15 min, then in TBS, blocked at 37 °C for 30 min, washed, and incubated with primary antibody at 4 °C overnight, followed by HRP-conjugated secondary antibody diluted in 5% BSA/TBS for 1 h at RT. DAB (Sigma, D8001) containing 1% $H_2O_2$ was applied as a chromogen. Staining was monitored under LM; reaction was stopped by washing with TBS. Samples were then dehydrated in a graded ethanol series, cleared with xylene, and mounted in Eukitt Quick-hardening mounting medium (Sigma).

**Light microscopy and image processing**

Images were acquired using a Zeiss LSM 5 Exciter confocal microscope (LSM) equipped with Imager.M1 and a NeoFluar objective lens (63x/NA 1.40). The laser lines used were 405, 488, 543, and 633 nm. Fluorescence images were acquired, some of them with additional differential interference contrast (DIC) overlay. The system was operated with Zen 2008 software (Zeiss). Immunoperoxidase, Periodic acid-Schiff (PAS), Masson´s trichrome, and Sirius red-stained sections were examined in bright-field microscopy under a Zeiss Axio Imager Z2 LM equipped with an ApoTome2 structured illumination acquisition system and a Plan-Apochromat 20x/0.8 objective using Zeiss ZEN 2012 software (blue edition). All image processing was done with ImageJ (NIH).

**Filtration Slit Density Analysis**

Kidney sections (2 µm) were deparaffinized and rehydrated, followed by boiling for 5 min in Tris-EDTA buffer (10 mmol/l Tris, 1 mmol/l EDTA, pH 9) in a pressure cooker for antigen retrieval. Next, sections were blocked in blocking solution (1% fetal bovine serum, 1% BSA, 0.1% fish gelatine, 1% normal goat serum) for 1 h. Primary antibodies (Suppl. Table S1) were incubated overnight at 4°C. After washing three times in PBS, secondary antibodies (Suppl. Table S1) were incubated for 1 h at RT. Nuclei were counterstained with DAPI for 5 min, followed by a washing step in PBS. Finally, the sections were washed in distilled $H_2O$ and mounted in Mowiol 4-88 (Carl Roth) using high-precision cover glasses (Paul Marienfeld). The evaluation of the filtratin slit density (FSD) was performed according to podocyte exact morphology measurement procedure (PEMP) [s1,2]. 3D-structured illumination microscopy (3D-SIM) was performed with Z-stacks from 19 planes with 488 and 561 nm channels acquired from the stained kidney sections using an N-SIM super-resolution microscope (Nikon) equipped with a 100x silicone objective. The Z-stacks were converted into a maximum intensity projection followed by the automatized identification of the filtration slit length as an index of foot process effacement. FSD was expressed as the ratio of the total filtration slit diaphragm per podocyte foot process area. FSD values of 20 glomeruli per animal were quantified in at least n=3 rats per group.

**Electron microscopy**

Samples from perfusion-fixed and post-fixed kidneys were embedded in Epon or transferred to critical-point drying for conventional TEM, large-scale scanning TEM (STEM), or SEM, respectively. Semithin sections (1 µm) were prepared from Epon blocks with an ultramicrotome (Ultracut E, Reichert-Jung) and stained with Richardson's stain for LM evaluation or to select a region for conventional or large-scale digitization. To screen pathological patterns ultrastructurally, conventional or large-scale ultrathin sections (200 to 400 nm) were prepared [s3]. Sections for large-scale analysis were partly or completely digitized at 3 to 4 nm pixel size using TrakEM2 for stitching and nip2 for export to high-resolution tif files and inspection with QuPath. Qualitative and quantitative analyses were performed using either a Gemini 300 field emission scanning electron microscope (FESEM, Zeiss) equipped with a scanning transmission electron microscopy (STEM) detector, SmartSEM, and Atlas 5 software. Alternatively, a Zeiss EM901 was used for conventional TEM. For repository implementation on www.nanotomy.org and open access pan-and-zoom analysis, datasets were exported into a tiled, browser-based file format using Atlas 5 software. For conventional 3D-SEM, dried and sputter-coated samples were evaluated by FESEM using an SEM detector

**Morphometric analysis**

To assess tubulo-interstitial fibrosis from PAS-stained paraffin sections, brightness and contrast were adjusted in ImageJ. Lines were drawn around the perimeter of regions of interest (ROI);

original magnification was 200x. Data were expressed as positively stained ROI vs. selected field areas. All samples were examined in a blinded manner. For quantitative assessment of alpha-smooth muscle actin (α-SMA) and renin immunoperoxidase staining, at least 15 non-overlapping fields were selected and α-SMA- and renin-immunoreactive ROI evaluated using ImageJ. The average ratio of ROIs to each microscopic field (200x magnification) was calculated and graphed. All measurements were performed by a single operator in a blinded fashion. Estimation of glomerular podocyte intactness was performed with WT1 and DAPI immunofluorescence staining using paraffin sections. Fluorescence signal was acquired on whole-slide images. Podocyte numbers per glomerulus were calculated from all glomeruli per section. Endothelial pore density was determined by counting the number of fenestrae per micrometer of glomerular basement membrane (GBM) in TEM images; total length of GBM was >750 µm per animal. At least 6 glomeruli per animal were evaluated. To determine fenestration density of cortical peritubular capillaries, TEM images were used. The number of fenestrae per micrometer of basement membrane length was counted. Ten cross-sectional profiles per animal were evaluated; total length of basement membrane was >850 µm per animal. Lysosome numbers in proximal tubules were counted on Richardson’s stained semithin plastic sections. Conventional lysosomes were termed “dark” based on their dark-blue stained core; those with clear-appearing cores were termed “heterolysosomes”. Twelve to 15 fields selected randomly, each containing 3 to 4 PCT profiles, were examined at 200x magnification. Wall-to-lumen ratios of renal afferent arterioles were analyzed in PAS-stained paraffin sections. Only profiles with open lumen were chosen. Wall and lumen surface areas were calculated using ImageJ by drawing lines around outer and inner perimeter. Five to 8 arterioles were studied per animal. The mean glomerular tuft area was measured by using anti-podocin-stained paraffin sections from entire kidneys imaged with the 3D-SIM. All available glomeruli with identifiable vascular pole were evaluated per section, their numbers ranging between 182 to 343. Podocin-immunoreactive area was quantified as $µm^2$ (NIPOKA). Quantification of anti-catalase immunofluorescence staining was performed by randomly selecting 10 optical fields, each containing 3 to 5 PCT profiles per slide at 200x magnification. Adjustments of pinhole, laser power, offset gain, and detector amplification below pixel saturation were maintained constant throughout. Mean fluorescence of catalase per proximal convoluted tubule (PCT) was determined with ImageJ. At least n=4 rats per group were evaluated throughout except for PAS-fibrosis and arteriolar morphometric measurements (n=3 rats).

**TUNEL assay**

Deparaffinized kidney sections (4 µm) were post-fixed in 4% PFA and labeled with TUNEL (Abcam, Ab66108) according to manufacturer's instructions in order to detect DNA fragmentation. Nuclei were counterstained with DAPI. Apoptotic signals were acquired by fluorescence microscopy (200x magnification) with DIC overlay. Labeled nuclei were counted in randomly chosen proximal tubular profiles from at least 10 adjacent optical fields and at least 15 randomly chosen glomeruli per animal, respectively. The apoptosis rate was calculated as the ratio of TUNEL-positive nuclei per total number of nuclei and field or glomerulus. At least n=4 rats per group were used for each experiment. Image analysis was performed with ImageJ.

**Western Blotting**

Whole kidney tissue was ground in liquid nitrogen and lysed in homogenization buffer (250 mM sucrose, 10 mM triethanolamine [AppliChem, PanReac, ITW Reagents]), supplemented with a protease inhibitor cocktail (cOmpleteTM, Roche), and sonicated 4 times for 1 s each. The supernatant of the protein lysates was obtained by centrifugation at 1.000xg for 10 min at +4 °C. Protein concentrations were measured using Micro BCA™ protein-assay-kit (Thermo Scientific). Samples were stored at -80 °C until further processing. Samples were then mixed 1:4 with 4x Laemmli buffer (Bio-Rad) containing 10% (v/v) β-mercaptoethanol (Merck), incubated at 65 °C for 10 min, separated by SDS PAGE (10% or 14%; 30 to 40 µg per lane), and transferred to a PVDF membrane (Macherey-Nagel). Membranes were then blocked with 5% milk or BSA in TBS and incubated with primary antibody overnight at +4 °C on a rotating wheel. Antibodies used for immunoblotting are listed in Suppl. Table S1. HRP-conjugated secondary antibody (Dako; diluted 1:2.000) was applied for 1 h at RT. Signal was generated by chemiluminescent reagent (Amersham ECL Western blotting detection reagent, GE Healthcare). Blots were imaged using an Intas ECL ChemoCam Imager (Intas Science Imaging). Densitometric quantification was performed using ImageJ.

**RNA extraction and qualification**

Whole kidney total RNA was isolated using PeqGOLD TriFast (VWR Life SCIENCE) according to manufacturer's instructions. The purity of RNA was checked using NanoPhotometer R spectrophotometer (IMPLEN). Next, an RNA Nano 6000 Assay Kit and Bioanalyzer 2100 system (Agilent Technologies) were used to evaluate quantity and integrity of RNA.

**RNA sequencing and data processing**

RNA-seq of rat kidney RNA samples (n=4 to 6 per group) was performed (Novogene, https://en.novogene.com/). A total amount of 1 μg RNA per sample was used as input material. Sequencing libraries were generated using NEBNext RNA Library Prep Kit for Illumina (New England BioLabs), and index codes were added to attribute sequences to each sample. Clustering of the index-coded samples was performed on a cBot Cluster Generation System with PE Cluster Kit cBot-HS (Illumina). After cluster generation, libraries were sequenced using Novaseq HisEquation 4000 platform (Illumina) and 150 bp paired-end reads were generated. Differential expression analysis between two groups was performed using DESeq2 R package. The resulting *P* values were adjusted using Benjamini and Hochberg's approach for controlling the False Discovery Rate (FDR). Genes with an adjusted *P* value < 0.1 (with no logFC cutoff) found by DESeq2 were considered as differentially expressed. RNA-seq data were deposited in the NCBI's Gene Expression Omnibus repository (GSE225215).

**Global proteomics**

Global proteomics was performed by using protein lysates of rat kidneys (n=4 to 6 per group) according to a previously established methodology [s4] (Charité Core Facility for High Throughput

Mass Spectrometry, Berlin). Briefly, protein lysates were trypsinized followed by analyzing tryptic peptides by LC-MS/MS using a timsTOF Pro 2 mass spectrometer (Bruker). The raw data were processed using DIA-NN 1.8 [s5] with standard settings using MS1 and MS2 resolution of 10 ppm. Peptides were identified by library-free mode using the *Rattus norvegicus* UniProt (UniProt Consortium 2019) sequence database (UP000002494_10116, downloaded on 20210116) and the matched-between-runs (MBR) option. The output was filtered at 0.01 FDR at the peptide level. All further analyses were performed using the R package DEP [s6]. Proteins with an adjusted $P$ value < 0.1 (with no logFC cutoff) were considered as DEP. Data are deposited in the ProteomeXchange Consortium via the PRIDE partner repository with an accession number: PXD038841.

**Global phosphoproteomics**

Global phosphoproteomic analyses were perfomed [s7] using cryo-pulverized rat kidney tissues (n=4 to 5 per group) (Proteomics/Max Delbrück Center, Berlin). Samples were lysed in SDC lysis buffer (1% Na-deoxycholate, 150 mM NaCl, 50 mM Tris-HCl pH 8.0, 1 mM EDTA, 10 mM DTT, 40 mM CAA [2-chloroacetamide, Sigma], phosphatase inhibitor cocktail II and III [Sigma]) by heating to 95 °C for 10 min. The solution was then cooled to RT, and Benzonase® (Merck; 50 units) was added for 30 min at 37 °C. Protein (200 µg per sample) was digested overnight at 37 °C with endopeptidase LysC (Wako) and sequence-grade trypsin (Promega) at a 1:100 enzyme-to-protein ratio. For relative quantification of phosphoproteins by liquid chromatography tandem mass spectrometry (LC-MS/MS), 200 µg per samples were labeled with TMTpro reagents (CsA sample set) or TMT10 (Tac sample set) according to manufacturer’s protocols (Thermo Fisher Scientific). Isobarically-labeled peptides were combined and fractionated into 30 fractions using high pH-reversed phase chromatography. Samples of each fraction underwent phosphopeptide enrichment using immobilized metal affinity chromatography (IMAC). Flow-throughs after IMAC enrichment were collected and dried. Resulting powders were dissolved and separated in a high performance liquid chromatography system (Thermo Fisher Scientific). Samples were measured in a Q-Exactive HF-X instrument (Thermo Fisher Scientific) operated in data-adapted acquisition mode. Raw data were analyzed using v 1.6.10.44 MaxQuant software package [s8]. The internal Andromeda search engine was used to search MS2 spectra against a decoy rat UniProt database (release 2019-07) containing forward and reverse sequences. The FDR was set to 1% for peptide and protein identifications, respectively. Unique and razor peptides were included for quantification. Statistical analysis was done with the Perseus software (version 1.6.2.1). DEPP were calculated using Student´s *t*-test with FDR based cutoff of 5%. The data presented in the study are deposited in the ProteomeXchange Consortium via the PRIDE partner repository, accession number PXD038546.

**Gene ontology and pathway analyses**

The Gene Ontology (GO) and pathway analyses for differentially expressed genes, proteins and phosphoproteins were performed using online bioinformatics tool Enrichr (https://maayanlab.cloud/Enrichr/). Pathways with a significance of $P < 0.05$ were defined as significantly regulated.

**Statistics**

Results were done in an observer-blinded way and analyzed using routine parametric statistics for normal distribution as assumed from the experimental design. Comparative analysis between two groups was performed by Student's *t*-test. Evaluation of multiple groups was performed using ANOVA followed by Tukey's post hoc test. GraphPad Prism7 software was used to analyze parameters. A significance level of $P < 0.05$ was accepted as significant.

**Supplementary Figures**

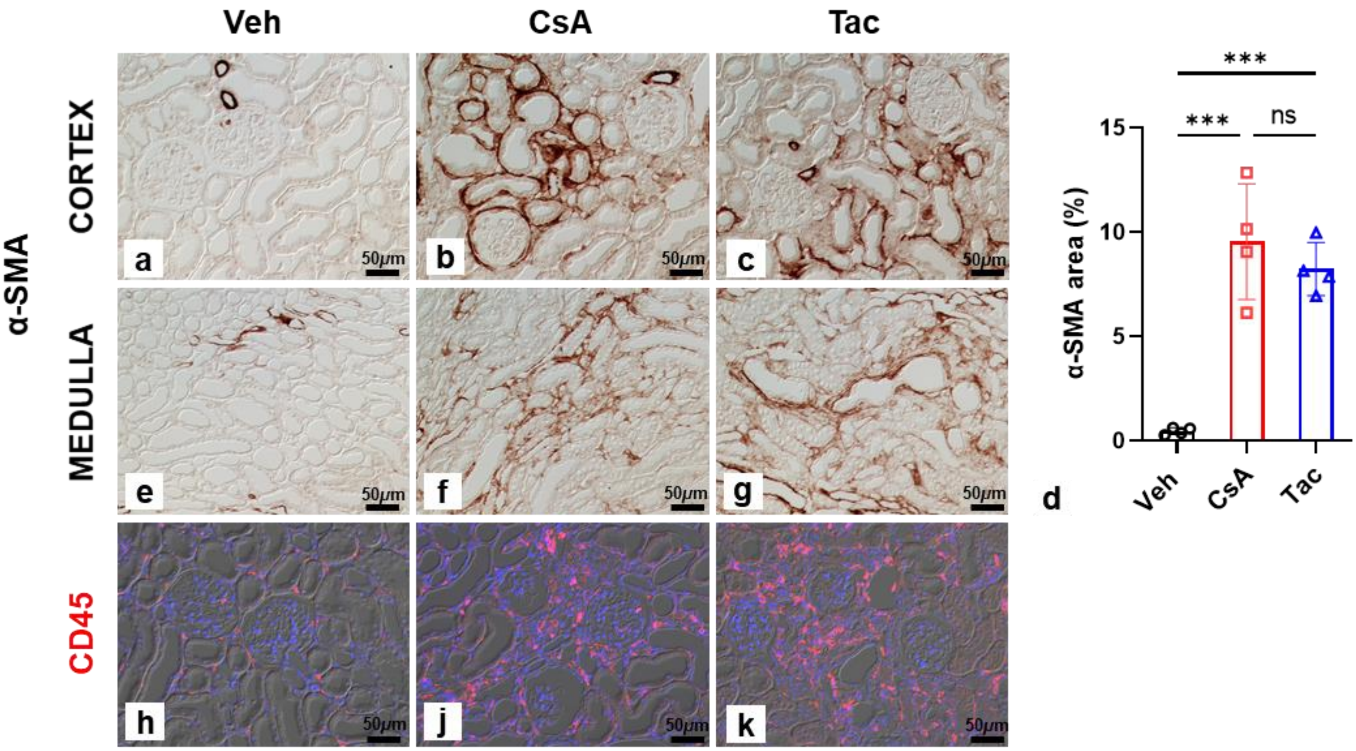


**Supplementary Figure S1. Smooth muscle actin (α-SMA) and leukocyte common antigen (CD45) staining**. (a-g) Anti-α-SMA immunoperoxidase staining in vehicle (Veh) (a,e) shows vascular wall signals. In cyclosporine A (CsA) and tacrolimus (Tac), interstitial signals are enhanced in foci around tubules and glomeruli, and in outer stripe focal areas. Signal is pronounced near atrophic or necrotic tubules and adjacent glomeruli, and pericapsular expression is variably present with stronger signals encountered in the CsA than in the Tac group (b, c, f, g). Bar graphs indicate average size of α-SMA-stained cortical foci (percent of unit sectional area; d); values are means ± SD; ****P*<0.001; ns, not significant. (h-k) CD45 immunofluorescence staining shows scattered individual interstitial cells in the cortex of Veh (h) and enhanced interstitial signals in fibrotic foci of CsA and Tac samples (j, k). DIC optics; bars indicate magnification. Statistical tests were performed using ANOVA with Tukey’s multiple comparison test (d).

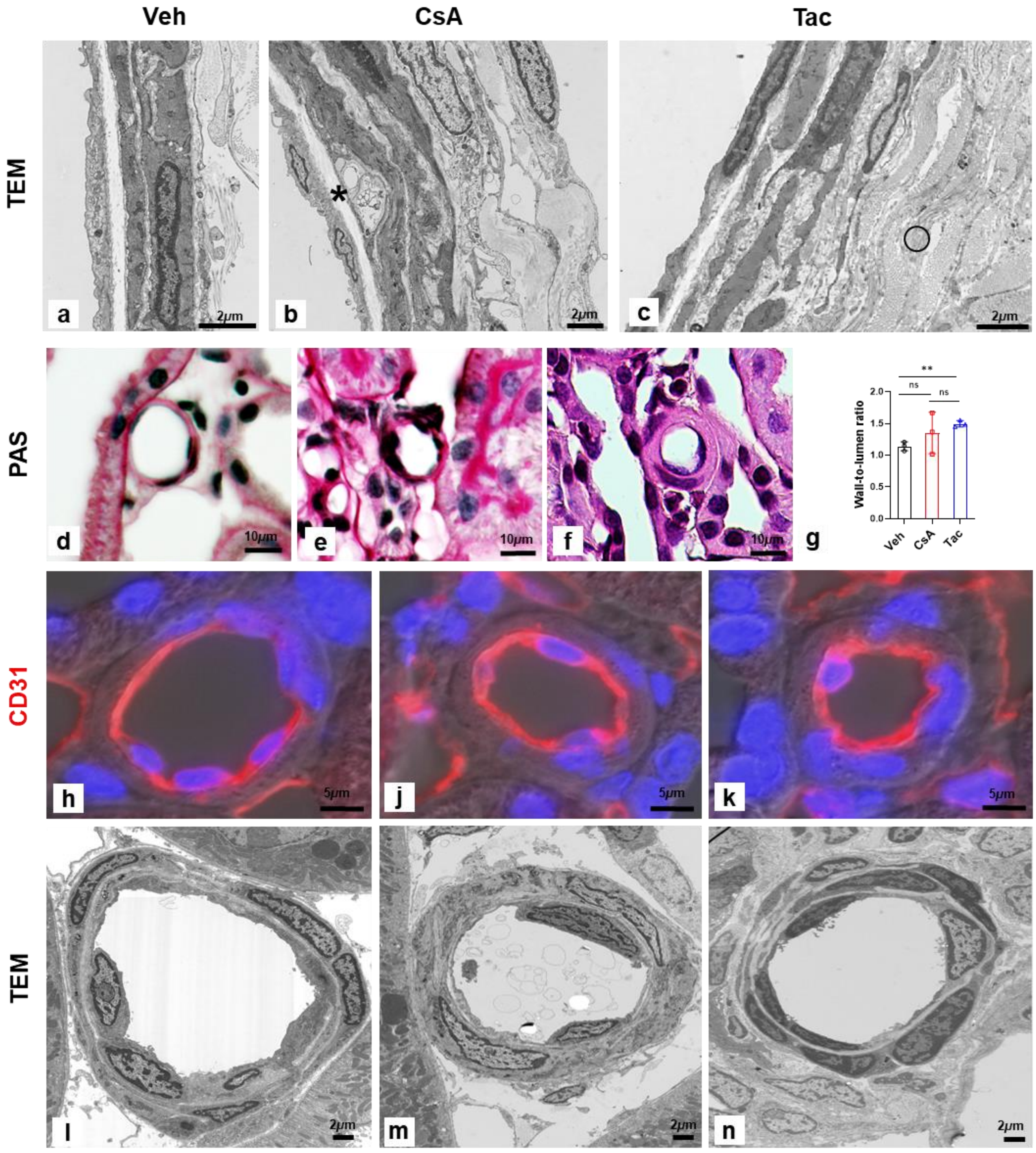


**Supplementary Figure S2. Renal arterial and arteriolar wall structure**. (a-c) Interlobular arterial walls are compared in representative views; in a vehicle (Veh) sample, endothelium, media, and adventitia present normal structure (a). In cyclosporine A (CsA), occasional vesicular or hyaline inclusions, here in subintimal location (asterisk in b) are shown. In tacrolimus (Tac), thinning of media wall cells, augmented matrix deposition, and thickened adventitia with substantial fibrillar collagen I layers (open circle) are characteristic (c). (d-g) Afferent arterioles, Periodic acid-Schiff (PAS); note the distinct staining intensities; luminal occlusion was generally not observed. (d-f). Wall-to-lumen ratios (g); values are means ± SD; ** $P<0.01$; ns, not significant. (h-k) Wall structure and endothelial CD31 immunoreactivity; note stronger signal in CNI groups. (l-n) Afferent arteriolar wall structure by TEM. DAPI blue nuclear staining and DIC optics (h-k); bars indicate magnification. Statistical tests were performed using ANOVA with Tukey's multiple comparison test (g).

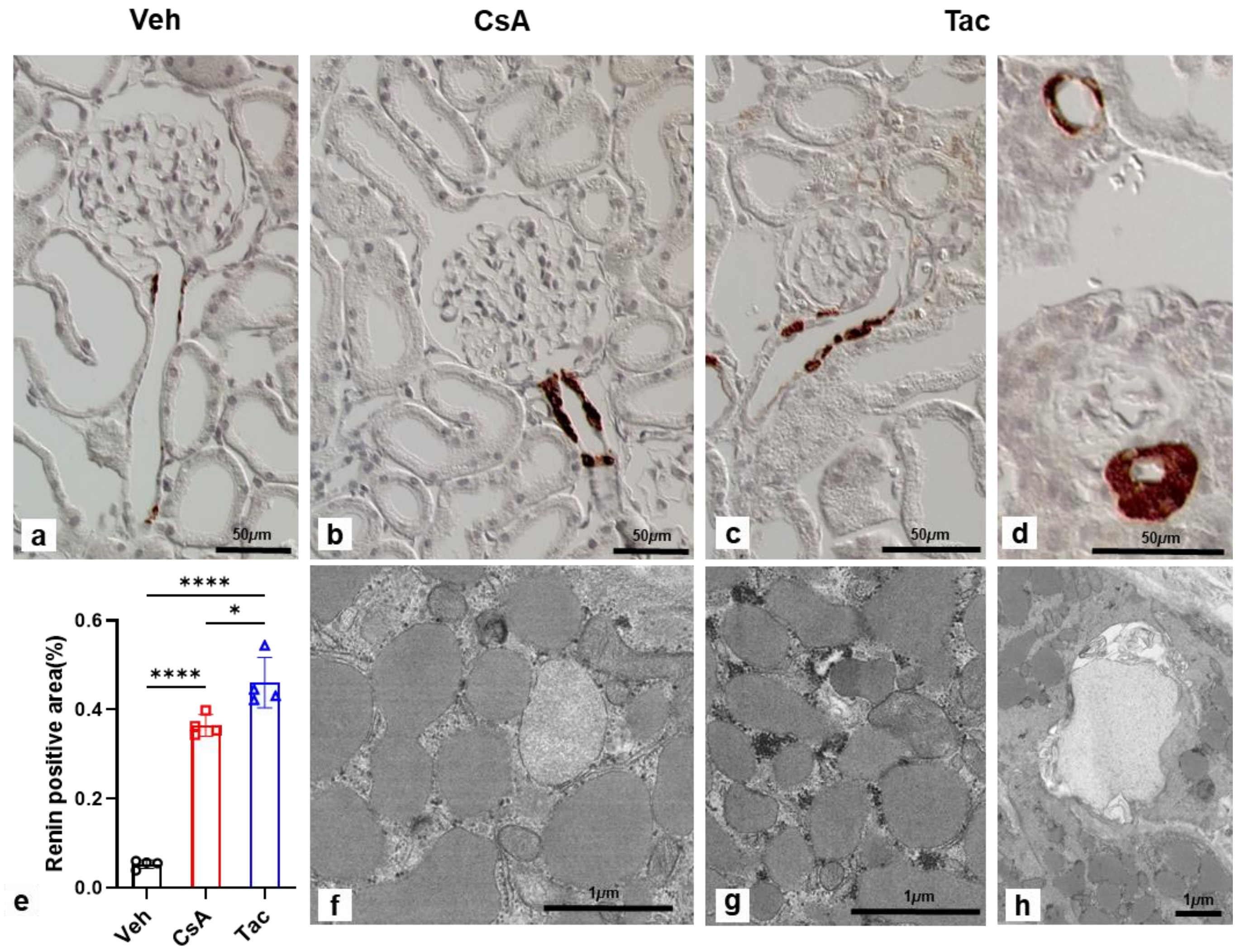


**Supplementary Figure S3. Juxtaglomerular apparatus – renin. (**a-d) Anti-renin immunoperoxidase staining shows normal distribution of renin signal in the preglomerular afferent arteriolar portion (a), highly increased signal in cyclosporine A (CsA) (b), and even higher increase in tacrolimus (Tac) (c) in representative views. In Tac, narrowing of the preglomerular afferent arteriolar portion of retracted-appearing glomeruli occurs (d); here, granular cells may form layers. (e) Bar graphs show quantitative expression of renin (immunoreactive area as percent of unit sectional area); values are means ±SD; * $P<0.05$, **** $P<0.0001$. (f-h) TEM shows high glycogen deposits in juxtaglomerular granular cells next to the renin granules in Tac (g) compared to CsA (f), and necrotic inclusions may be observed in granular cell fields in Tac (h). DIC optics, hematoxylin stain (a-d); bars indicate magnification. Statistical tests were performed using ANOVA with Tukey's multiple comparison test (e).

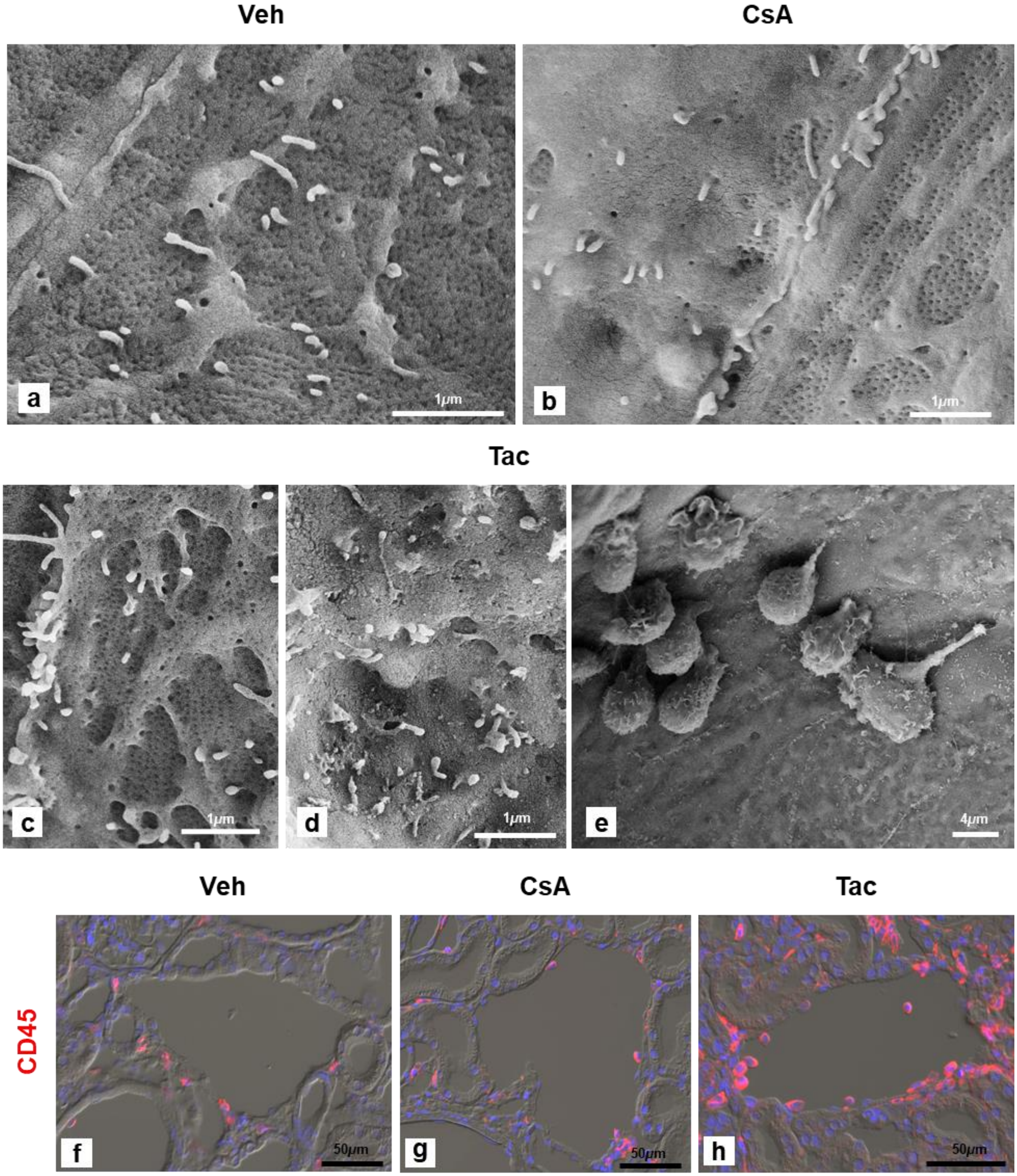


**Supplementary Figure S4. Renal venous endothelia.** (a-e) Representative SEM views of venous endothelia (arcuate or interlobular veins); typical distribution of fenestrated pore fields in a vehicle (Veh) sample (a); similar structure is seen in cyclosporine A (CsA) next to a pore-free field (b), whereas in tacrolimus (Tac), normal pore fields (c) or larger areas of absent fenestration may be encountered (d); adherent leukocytes are frequent (e). (f-h) CD45 immunoreactivity shows scattered perivascular leukocytes in Veh (f), few adherent leukocytes in CsA (g), and numerous adherent leukocytes in Tac (h). DAPI blue nuclear staining and DIC optics (f-h). Bars indicate magnification.

.

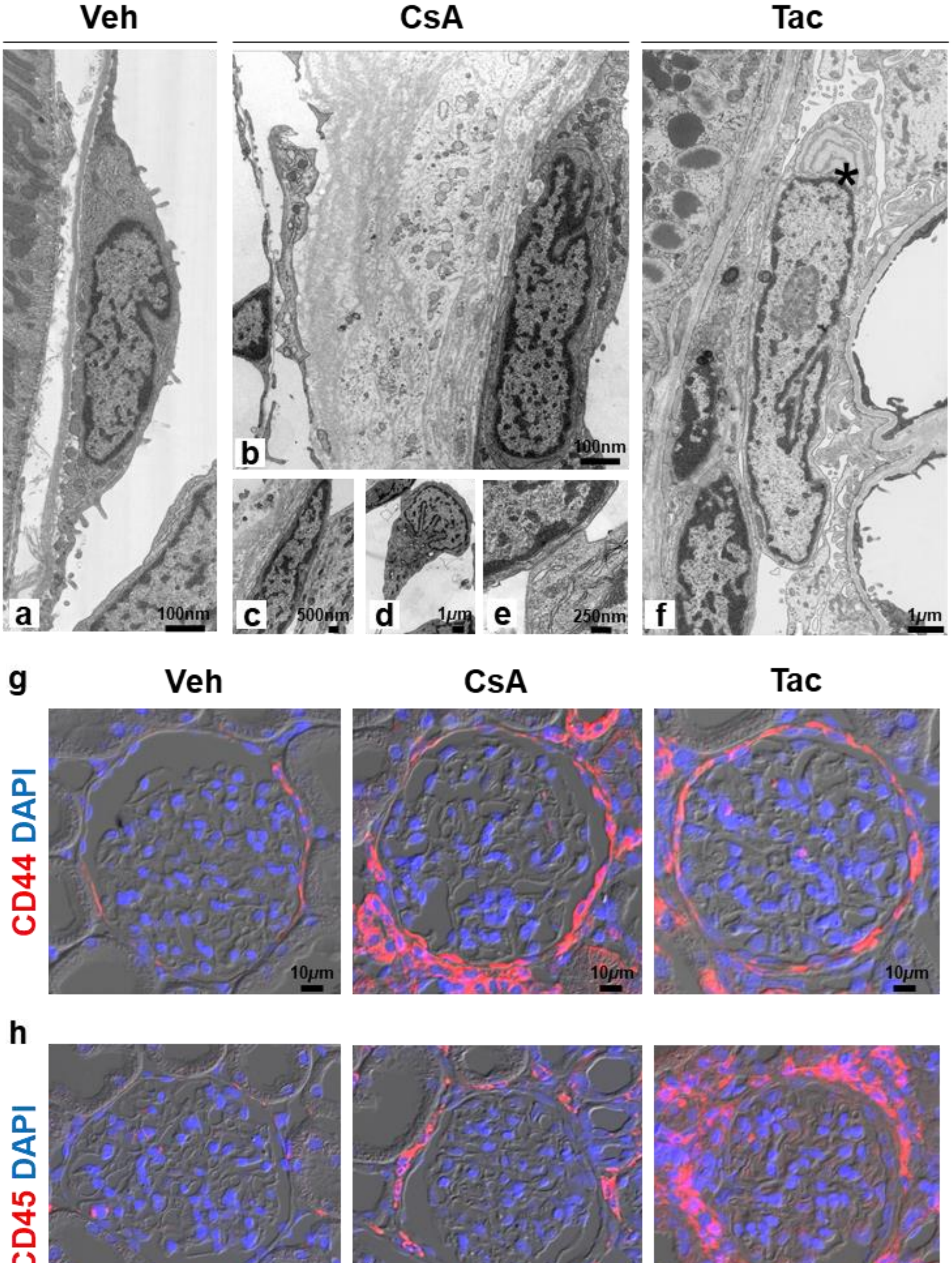

Veh
CsA
Tac
a
100nm
b
100nm
c
500nm
d
1μm
e
250nm
f
1μm
*
g
Veh
CsA
Tac
CD44 DAPI
10μm
h
CD45 DAPI
10μm

**Supplementary Figure S5. Changes in Bowman's capsule.** (a-f) TEM representative images show regular structure of Bowman's capsule in vehicle (Veh) with flat parietal epithelium and thin capsular basement membrane (a). In cyclosporine A (CsA), both parietal epithelium and capsular basement membrane are markedly thickened, ranging from 0.2 up to 4 µm in cell height, with extensive granular and hyaline matrix formation (b), and synechiae with multiple contact points are formed (c); podocytes may form synechiae even in the urinary pole area where they may display tip lesions (d,e). In tacrolimus (Tac) a similar synechia between podocyte and parietal epithelium is shown; note typical rER inclusion body within podocyte (asterisk) (f). (g) Anti-CD44 immunofluorescence staining shows occasional mild signal in Bowman's capsule in a Veh sample, whereas in CsA and in Tac activated parietal epithelium presents with high signal intensity along almost the entire perimeter. (h) Anti-CD45 staining shows few interstitial signals in Veh, but strong signal associated with the capsule in CsA and, more so, in Tac. DAPI blue nuclear staining and DIC optics; bars indicate magnification.

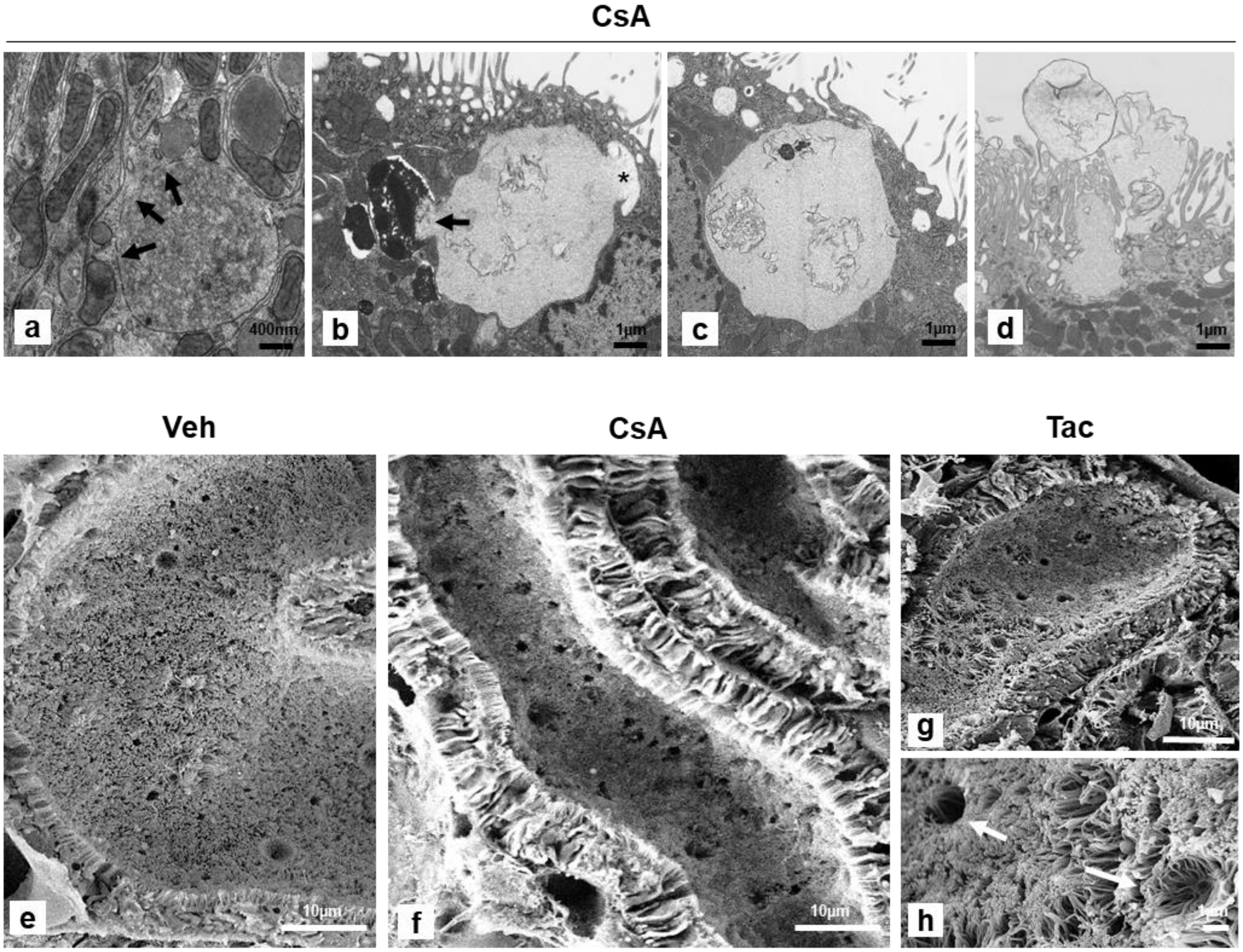


**Supplementary Figure S6. Ultrastructure of lysosomal canges in proximal tubule**

(a-d) TEM reveals that in cyclosporine A (CsA), heterolysosomes reveal partial communication with the cytosol via interrupted membrane portions (arrows; a), anastomose with residual bodies (arrow) and late endosomes (asterisk; b), are filled with heterogenous debris, reach the luminal cell membrane (c), and are frequently encountered in exocytotic transition (d). (e-h) Representative SEM views show the frequency of lysosomal exocytosis in proximal convoluted tubule by the number of transitory states, leaving pits in the brush border. Vehicle (Veh) defines is baseline activity (e), whereas in CsA exocytosis is substantially enhanced (f); in tacrolimus (Tac) there is intermediate activity (g); detailed view of pit formations (arrows; h). Bars indicate magnification.

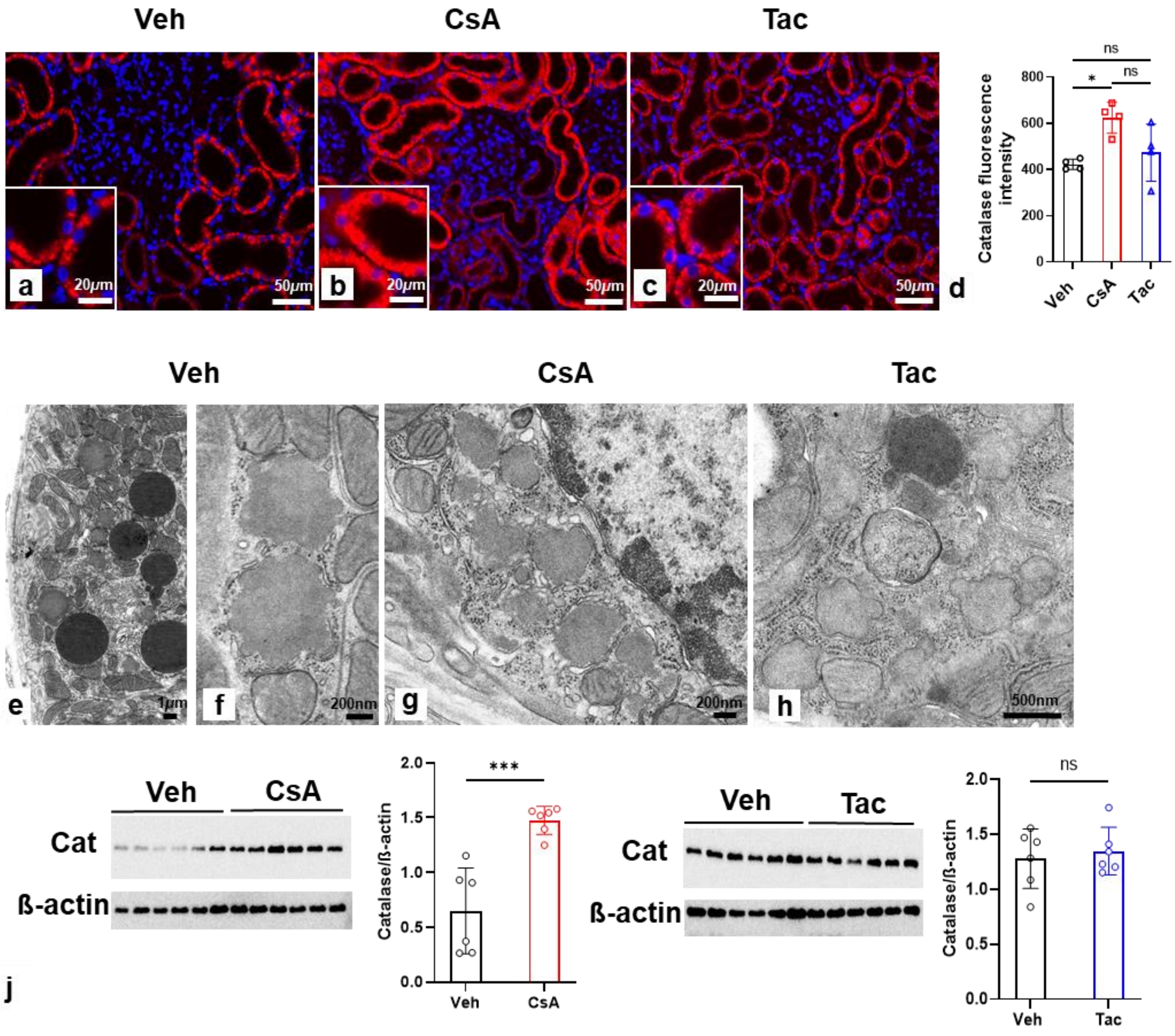


**Supplementary Figure S7. Catalase abundance and peroxisome structure in the proximal tubule.** (a-d) Immunofluorescence staining with anti-catalase antibody. There is abundant staining in proximal convoluted tubule of Vehicle (Veh) (a), cyclosporine A (CsA) (b), and tacrolimus (Tac) samples (c). Inserts reveal punctuate nature of the signal; DAPI blue nuclear staining. Note intensified signal in CsA. Bar diagram shows enhanced fluorescence signal quantified per unit sectional area from catalase-immunostained sections; values are means ± SD; * $P<0.05$; ns, not significant. (e-h) TEM images showing regular distribution of peroxisomes at the basal cell pole (e) and regular peroxisome morphology with adjacent cisternae of rough and smooth endoplasmic reticulum at higher resolution in Veh (f). Clusters of peroxisomes are visualized in CsA (g) and Tac (h); their extent may vary with treatment and cell affection.
(j) Western blots show immunoreactive catalase (60 kDa) from kidney extracts of CsA, Tac, and their respective vehicle groups (n=6, respectively); ß-actin (42 kDa) serves as loading control.

Densitometric evaluation is presented on the right side of the respective blots. Data are means ± SD; *** $P<0.001$, ns, not significant. Statistical tests were performed using ANOVA with Tukey's multiple comparison test (d) or unpaired Student's *t* test (j). Cat, catalase.

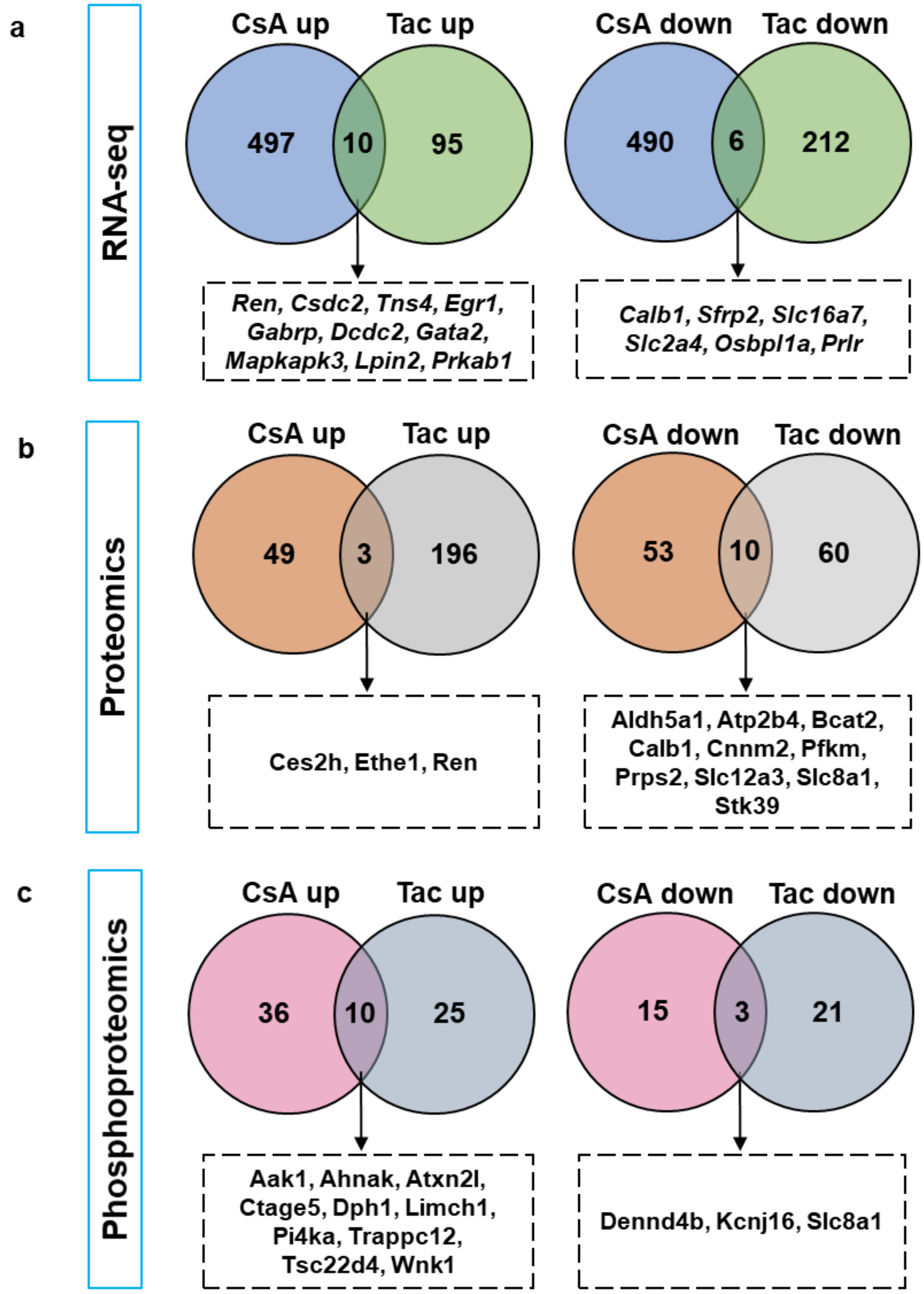


**Supplementary Figure S8. Differentially expressed genes, proteins, and phosphoproteins.** (a-c) Venn diagram showing the number of differentially expressed genes, proteins, and phosphoproteins (RNA-seq and proteomics, adjusted *P*< 0.1; phosphoproteomics, FDR<5; CNI vs. Veh). Jointly regulated products arelisted in box. “up”, upregulated; “down”, downregulated.

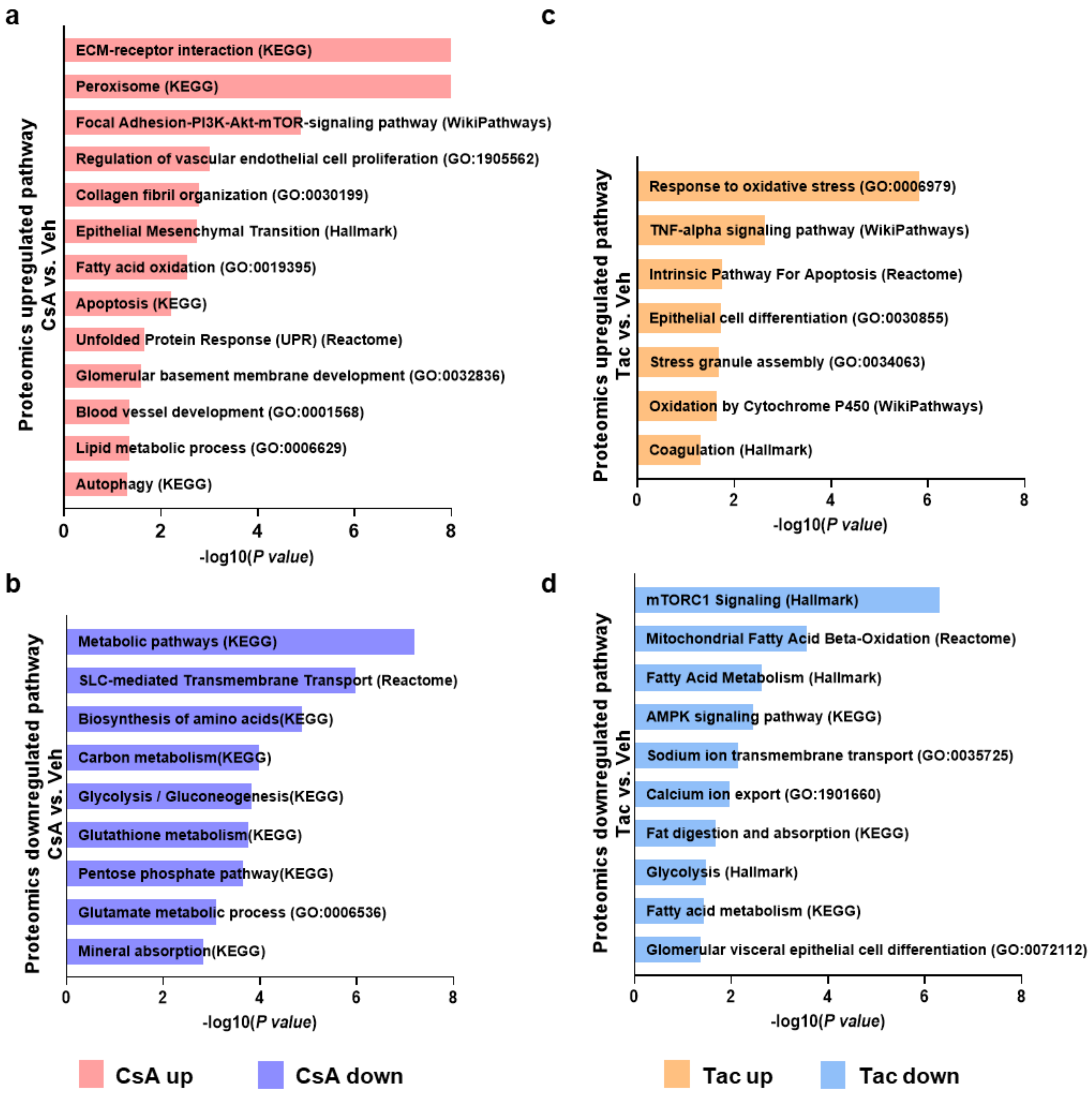


**Supplementary Figure S9. Pathway enrichment analysis of differentially expressed proteins (DEPs) affected by calcineurin inhibitor treatments.** Significantly enriched pathways in proteins up- (a) and downregulated (b) in cyclosporine A (CsA) versus vehicle (Veh) ($P < 0.05$). Significantly enriched pathways in proteins up- (c) and downregulated (d) in tacrolimus (Tac) versus vehicle (Veh) ($P < 0.05$). Term lists used in this analysis were GO_Biological_Processes, WikiPathways, Reactome, KEGG, and Hallmark to determine enriched processes and pathways from DEPs (Enrichr webtool).

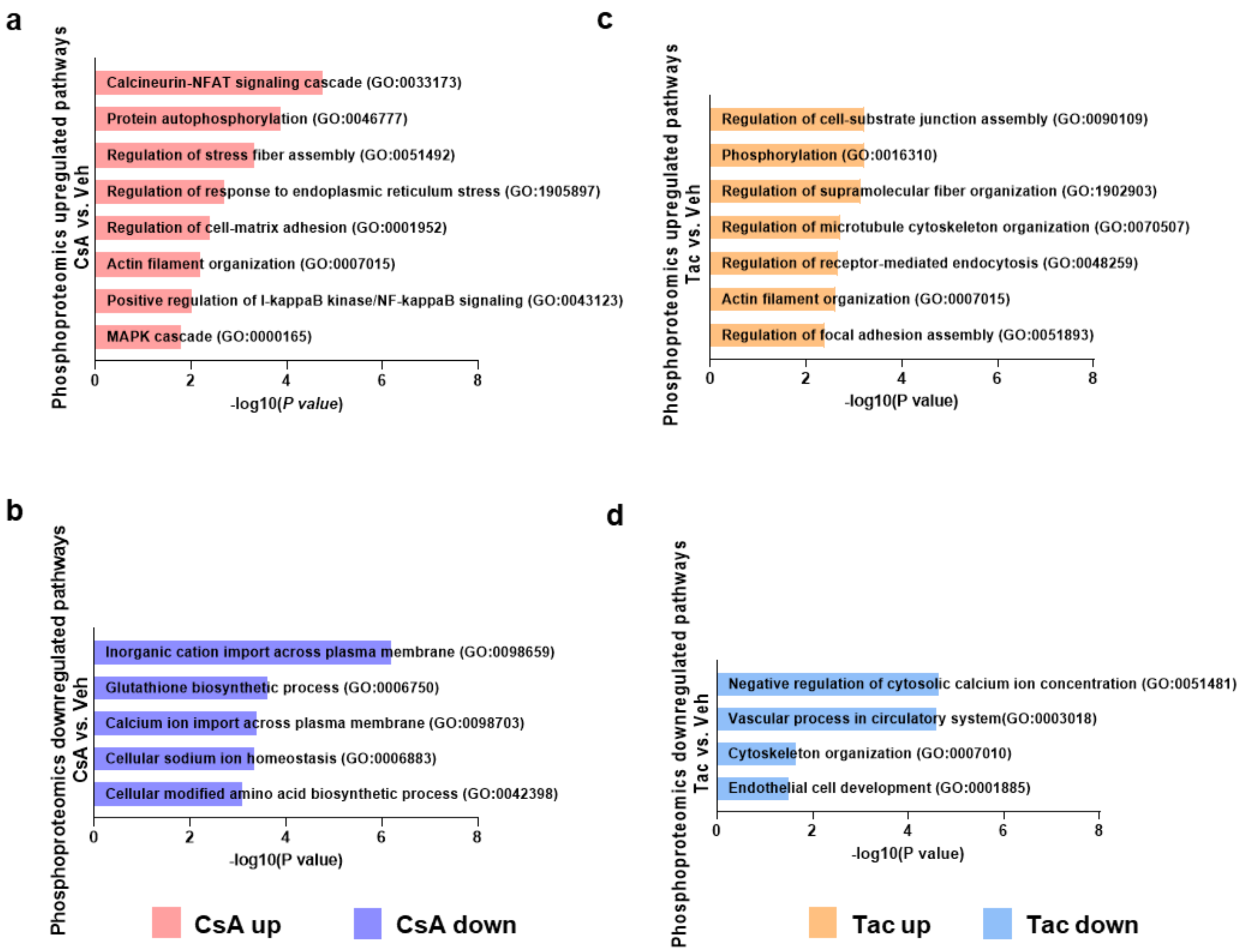


**Supplementary Figure S10. Pathway enrichment analysis of differentially expressed phosphoproteins (DEPPs) affected by calcineurin inhibitor treatments.** Significantly enriched pathways in phosphoproteins up- (a) and downregulated (b) in cyclosporine A (CsA) versus vehicle (Veh) ($P < 0.05$). Significantly enriched pathways in phosphoproteins upregulated (c) and downregulated (d) in tacrolimus (Tac) versus vehicle (Veh) ($P < 0.05$). Term lists used in this analysis were GO_Biological_Processes, WikiPathways, Reactome, KEGG, and Hallmark to determine enriched processes and pathways from DEPPs (Enrichr webtool).

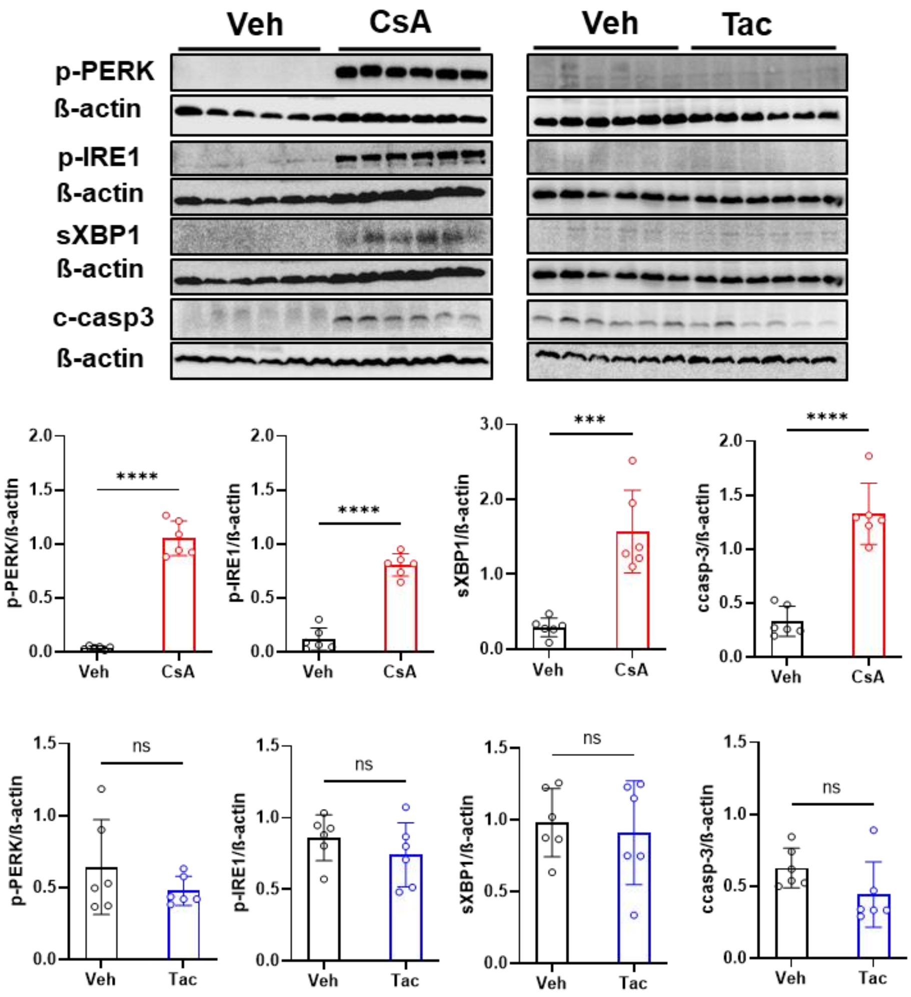


**Supplementary Figure S11. Western blot verification of unfolded protein response parameters**. Phospho-PKR-like ER kinase (p-PERK,Thr980,150 kDa), phospho- Inositol requiring-1α (p-IRE1α , Ser724, 110 kDa), X-box binding protein 1 (sXBP1) (55 kDa), cleaved caspase 3 (c-casp3) (17 kDa), and ß-actin (42 kDa), rat kidney. Below, corresponding densitometric evaluations expressed as –fold changes relative to vehicle (Veh) levels. Data are means ± SD; *n*=6 per group. ***$P < 0.001$, ****$P < 0.0001$; ns, not significant. Statistical tests were performed using unpaired Student’s t test.

**Differentially expressed PT marker genes / Csa vs. Veh**

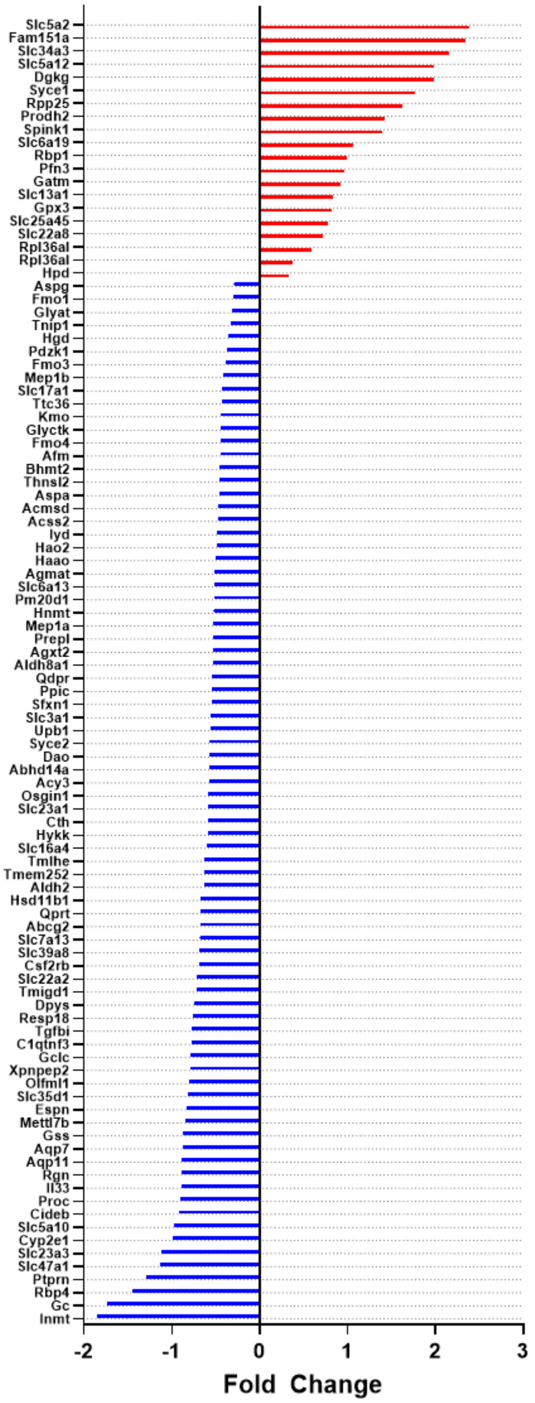


| Down | Up |
|---|---|
| 80 | 20 |

**Differentially expressed PT marker genes / Tac vs. Veh**

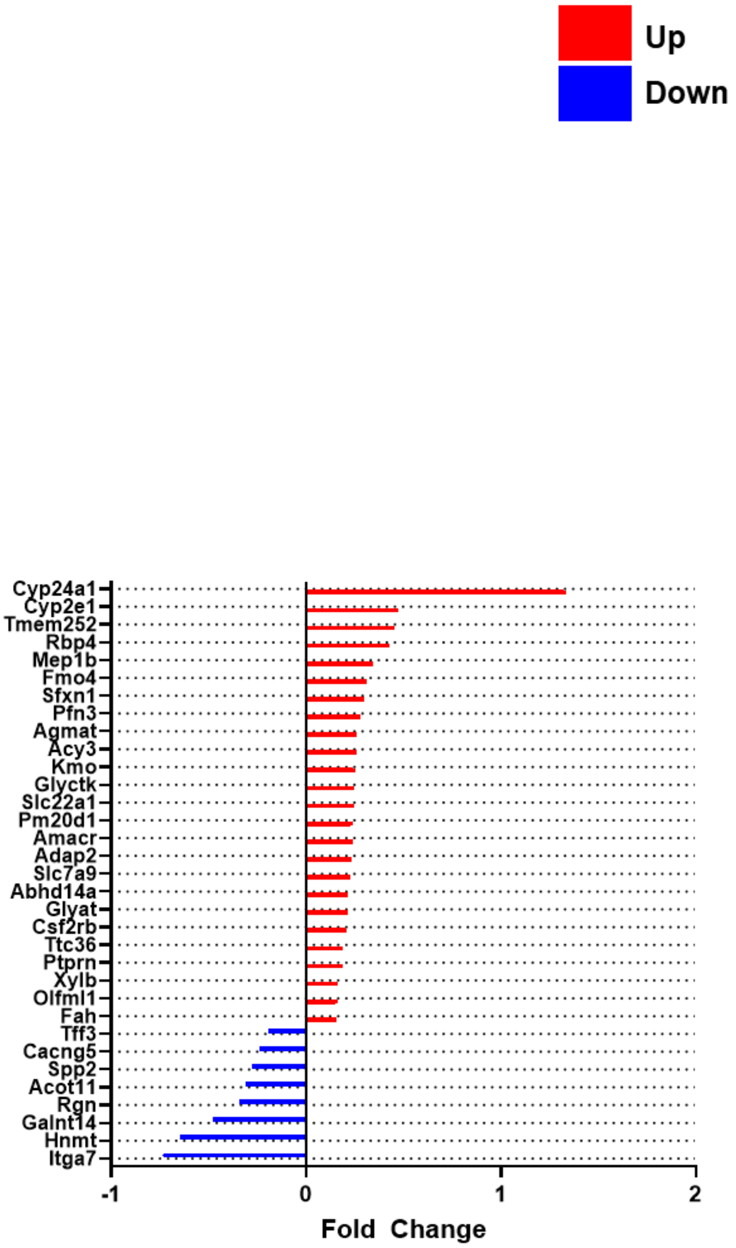


| Down | Up |
|---|---|
| 8 | 25 |

**Supplementary Figure S12. Differentially expressed genes with endogenous expression in proximal tubule.** Bar graph on a published RNA-seq data set of segmental profiles of the adult rat nephron [s9] showing 193 genes endogenously expressed in proximal tubule; 100 of these are differentially expressed in the cycslosporine A (CsA) group but only 33 in tacrolimus (Tac) ($P$ value < 0.05). Here, a tenfold higher number of transcriptionally downregulated genes in CsA compared to Tac should correspond to significant loss of function in this nephron segment. They indicate a tendency towards fibrosis and chronic kidney disease in CsA, such as upregulated *Sglt2* pointing to profibrotic changes of the S1 and S2 segments [s10], as well as downregulated *Hao2* standing for decreased fatty acid metabolism [s11], *Qdpr* for TGFß1, NOX1 and NOX4 stimulation [s12], *Tmigd1* for impaired protection from oxidative cell injury [s13], *C1qtnf3* for TGFß1 and interstitial fibrosis [s14], and *Kmo* for epithelial injury and allograft rejection [s15]. By contrast, in Tac stimulated *Cyp2e1* indicated activated resistance against epithelial decay which in this case may be reactive upon upstream glomerular injury [s16]. Veh, vehicle.

### Supplementary Tables

**Supplementary Table S1. List of antibodies used for immunohistochemistry and Western blot.** Listed are information on specification, host and clonality, source, dilution, and diluent

| Primary antibodies | Host & Clonality | Company | Catalogue Nr | IHC | WB |
|---|---|---|---|---|---|
| Anti-Cleaved Caspase 3 (Asp175) | rabbit, polyclonal | Cell Signaling Technology | 9661 | | 1:500, 5% BSA in TBS |
| Anti-Phospho-PERK (Thr980) | rabbit, monoclonal | Cell Signaling Technology | 3179 | | 1:1000, 5% Milk in TBST |
| Anti-Phospho-IRE1α (phospho S724) | rabbit, polyclonal | Abcam | ab48187 | | 1:1000, 5% BSA in TBST |
| Anti-ß-Actin | mouse, monoclonal | Sigma | A2228 | | 1:20000, 5% BSA in TBST |
| Anti-sXBP1 | rabbit, monoclonal | Abcam | ab220783 | | 1:1000, 5% BSA in PBS |
| Anti-LAMP1 | rabbit, polyclonal | Abcam | ab24170 | 1:2000, 5% BSA in TBS | |
| Anti-Renin | sheep, polyclonal | Acris | AP00945PU-N | 1:1000, 5% BSA in TBS | 1:1000, 5% BSA in TBS |
| Anti-CD31 | goat, polyclonal | R&D Systems | AF3628 | 1:500, 5% BSA in TBS | |
| Anti-CD44 | rabbit, monoclonal | Abcam | ab189524 | 1:2000, 5% BSA in TBS | |
| Anti-CD45 | Rabbit, polyclonal | Abcam | ab10558 | 1:1000, 5% BSA in TBS | |
| Anti-Catalase | rabbit, polyclonal | Abcam | ab217793 | 1:3000, 5% BSA in TBS | 1:2000, 5% BSA in TBS |
| Anti-α-sma | mouse, monoclonal | Sigma-Aldrich | A2547 | 1:1000, 5% BSA in PBS | 1:2000, 5% BSA in TBS |
| Anti-Wilms Tumor Protein-1 | rabbit, monoclonal | Abcam | ab89901 | 1:500, 5% BSA in TBS | |
| Anti-Integrin alpha 3 | goat, polyclonal | Abcam | ab223661 | 1:150,1% FBS, 1% BSA, 0.1% fish gelatine, 1% normal goat serum | |
| Anti-Podocin | rabbit, polyclonal | Immuno-Biological Laboratories | 29040 | 1:150,1% FBS, 1% BSA, 0.1% fish gelatine, 1% normal goat serum | |
| **Secondary antibodies** | **Host & Clonality** | **Company** | **Catalogue Nr** | | |
| Cy3-coupled donkey anti-rabbit IgG | Donkey, polyclonal | Dianova | 711-165-152 | | |
| Cy2-coupled donkey anti-rabbit IgG | Donkey, polyclonal | Dianova | 711-225-152 | | |
| Cy3-coupled donkey anti-mouse IgG | Donkey, polyclonal | Dianova | 715-165-150 | | |
| Cy2-coupled donkey anti-mouse IgG | Donkey, polyclonal | Dianova | 715-225-150 | | |
| Cy3-coupled donkey anti-goat IgG | Donkey, polyclonal | Dianova | 705-165-147 | | |
| Cy2-coupled donkey anti-goat IgG | Donkey, polyclonal | Dianova | 705-225-147 | | |

| | | |
|---|---|---|
| Swine Anti-Rabbit Immunoglobulins HRP | Dako | P0217 |
| Goat anti-mouse Immunglobulins/HRP | Dako | P0447 |
| Polyclonal Rabbit Anti-Goat Immunoglobulins/HRP | Dako | P044901-2 |
